# On the estimation of genome-average recombination rates

**DOI:** 10.1101/2023.09.01.555840

**Authors:** Julien Y. Dutheil

## Abstract

The rate at which recombination events occur in a population is an indicator of its effective population size and the organism’s reproduction mode. It determines the extent of linkage disequilibrium along the genome and, thereby, the efficacy of both purifying and positive selection. The population recombination rate can be inferred using models of genome evolution in populations. Classical methods based on the patterns of linkage-disequilibrium provide the most accurate estimates, providing large sample sizes are used and the demography of the population is properly accounted for. Here, the capacity of approaches based on the sequentially Markov coalescent (SMC) to infer the genome-average recombination rate from as little as a single diploid genome is examined. SMC approaches provide highly accurate estimates even in the presence of changing population sizes, providing that (1) within genome heterogeneity is accounted for and (2) classical maximum-likelihood optimization algorithms are employed to fit the model. SMC-based estimates proved sensitive to the presence of gene conversion, which leads to an overestimation of the recombination rate if conversion events are frequent. Conversely, methods based on the correlation of heterozygosity succeed in disentangling the rate of crossing over from that of gene conversion events, but only when the population size is constant and the recombination landscape homogeneous. These results call for a convergence of these two methods to obtain accurate and comparable estimates of recombination rates between populations.

## 1 Introduction

Recombination is a fundamental process impacting the segregation of alleles in populations and, consequently, a driver of genetic diversity between populations and within genomes [Stapley et al., 2017]. The amount of recombination occurring in a population is an indicator of the species’ reproduction regime, but also of the efficacy of selection within populations. Both aspects have implications for biodiversity characterisation (e.g. plankton species Rengefors et al. [2017]), species conservation [Theissinger et al., 2023], as well breeding strategies [Epstein et al., 2023].

Recombination rates can be inferred with different methods involving various degrees of laboratory work [Peñalba and Wolf, 2020]. The most experimentally accessible methods involve genome (re)sequencing, in combination with variant calling pipelines. These methods rely on the patterns that recombination leaves on the distribution of variants within populations; they infer the so-called population recombination rate, noted *ρ*, equal to 2 *· x · Ne · r*, where *Ne* is the effective population size, *x* the ploidy of the individuals serving as unit count for *Ne*, and *r* is the “molecular” recombination rate, the number of recombination events per base pair per generation.

The main signature of recombination in the sequence data used to infer recombination rates is linkage disequilibrium (LD) between variants. *LDhat* was a pioneering method for the reconstruction of fine-scale recombination maps from population genomics data [McVean et al., 2004]. The resulting inferred rate is population- and sex-averaged over time and individuals. Yet this method unravelled the variation of this rate along genomes with an unprecedented resolution. *LDhat* ‘s methodology served as the basis for two decades of developments, accounting notably for populations’ demography [Kamm et al., 2016] and improving scalability to increasingly larger data sets [Spence and Song, 2019].

In parallel to the development of methods to infer recombination rates, models based on the sequentially Markov coalescent (SMC) unleash the power of full genome data for demography inference [Spence et al., 2018]. The SMC is an approximation of the sequential coalescent with recombination [Wiuf and Hein, 1999]. While it makes simplifying assumptions upon the recombination process, SMC-based inference relies on the recombination rate, which can be jointly estimated with demographic parameters [Li and Durbin, 2011, Schiffels and Wang, 2020]. This property has been further used in the *iSMC* method, which accounts for the heterogeneity of the recombination rate assuming a prior distribution and an auto-correlation process along the genome [Barroso et al., 2019]. While SMC-based approaches do not reach a fine-scale resolution near that of LD-based methods, they permit the inference of recombination rates with as little as one unphased diploid genome.

Other approaches made use of LD properties to extract information about the recombination rate with small sample sizes. Just like SMC-based methods, methods relying on the correlation of heterozygosity provide estimates of genome-average recombination rates [Haubold et al., 2010]. However, such methods assume a homogeneous recombination rate along the analysed sequences and do not permit the inference of local variation. However, a recent extension, *heRho*, permits disentangling the crossing over (CO) rate from the rate of gene conversion (GC) events [Setter et al., 2022]. The performance of other LD-based and SMC-based methods in the presence of GC is yet to be characterised.

We here compare the performance of distinct population genomics methods to infer the genome average population recombination rate. This assessment is made with respect to sample size, the shape of the recombination landscape, the demography of the population, and the presence of gene conversion in addition to crossing-over events.

## 2 Materials and methods

### 2.1 Simulations

Simulated data sets of 10 Mb were generated using the *msprime* package version 1.2.0 [Baumdicker et al., 2022]. All simulations were performed with a per-generation mutation rate of 1.25e-8 bp*^−^*^1^ and an effective population size of 100,000 diploid individuals. Ten replicates were generated in each scenario. The resulting data sets were exported in the variant call format (VCF), the resulting VCF files serving as input for the various inference methods. The analysed scenarios varied according to the considered recombination landscape and demographic history. Several average recombination rates were considered: 0, 0.1, 0.5, 1, and 5 times 1.5e-8 M.bp*^−^*^1^, corresponding to *rho* = 4*.Ne.r* = 0, 0.0006, 0.003, 0.006, 0.03.

#### 2.1.1 Variable recombination landscape

Random recombination maps were generated by drawing segments with lengths taken from a geometric distribution of mean 10,000 bp. The recombination rate in each segment was drawn randomly from an exponential distribution with a mean of 1.0. The generated recombination maps were then multiplied by the average recombination rate.

#### 2.1.2 deCODE recombination landscape

As an alternative recombination map, the sex-averaged deCODE recombination map was used [Kong et al., 2002]. The first 10,000 10 kb windows of chromosome 1 were standardized in order to have a mean of 1.0 before being multiplied by the specified averaged recombination rate.

#### 2.1.3 Recombination landscape with hotspots

Hotspots were simulated by randomly sampling regions with lengths taken uniformly between 0.5 and 2 kb, for which a recombination rate of 70 *×* 1.25e-8 bp*^−^*^1^ was set. The distance between hotspots was taken from a geometric distribution with a mean of 50 kb. The recombination rate was uniform between two consecutive hotspots but different between inter-hotspot regions and was taken from an exponential distribution of mean 1.0. The “background” rate of the inter-hotspots region was then multiplied by the specified average recombination rate.

#### 2.1.4 Non-constant demographic scenarios

We considered two scenarios departing from a constant population size. In the first scenario (referred to as “decreasing population size” in the following), an ancestral population size of *Ne* = 100, 000 exponentially decreases to *Ne* = 1, 000 at present time, starting 1,000 generations before present. In a second scenario, the asymmetric scenario is considered where an ancestral population size of *Ne* = 1, 000 exponentially increases to *Ne* = 100, 000 at present, starting 1,000 generations before present. In this scenario, the average pairwise heterozygosity is 15% that of the constant or decreasing population size scenarios. We further considered a similar scenario where an ancestral population of 50,000 increased exponentially to a size of 200,000, which led to a genetic diversity of roughly half that of the constant or decreasing population size scenarios. When plotting the ‘true’ population recombination rate, the average *Ne* is estimated using the formula *Ne* = *π/*(4 *· u*), where *π* is the average pairwise heterozygosity of the simulated dataset (computed using the *vcftools*) and *u* = 1.25*e −* 8 is the mutation rate.

### 2.2 Inference of genome-average population recombination rate

For each simulated scenario – a combination of a recombination landscape, demographic scenario, average recombination rate, and sample size – we inferred the genome-average population recombination rate with distinct methods for ten independent replicates.

#### 2.2.1 The multiple sequentially Markov coalescent, version 2 (MSMC2)

Individual genome data were extracted from the simulated VCF files using the *vcftools* [Danecek et al., 2011], and then converted into *MSMC* input files using the provided python script generate_multihetsep.py. *MSMC2* [Schiffels and Wang, 2020] was run with default parameters and with a restricted epoch scheme with only 16 intervals, using the argument -p 1*2+11*1+1*3. Under the homogeneous rate scenario, *MSMC2* was also run with 40 iterations instead of the default 20 (argument -i 40), and with an initial *ρ/θ* = 4 instead of the default 0.5 (argument -r 4). The resulting population recombination rate was read from the last line of the output *.loop.txt file. If the program failed to converge and stopped before the 20 (resp. 40) iterations, the resulting estimate was recorded as missing data.

#### 2.2.2 The integrative sequentially Markov coalescent (iSMC)

*iSMC* [Barroso et al., 2019] was run on the simulated VCF files with a two-knots spline model, 40 time-intervals, and a precision of 0.1 on the log-likelihood. The method was referred to as *iSMC* when a homogeneous model was considered. We further considered a model with a five-class discrete gamma model of recombination, which we here note *rhoSMC*. Under this model, it is possible to use an empirical Bayesian approach to estimate the site-specific recombination by computing the mean of each recombination class weighted by their posterior probabilities, computed using the backward algorithm [Dutheil, 2021]. The posterior genome-average recombination rate was computed as the mean over all site-specific posterior estimates.

#### 2.2.3 LDhat

The simulated VCF files were converted to *LDhat* input files using the *vcftools* [Danecek et al., 2011]. No filtering on the minimum allele frequency was performed, and all SNPs were kept. The interval program from the *LDhat* package was run with options -its 10000000 -bpen 5 -samp 5000 (10 million iterations sampled every 5,000 generations, with a penalty of 5), as suggested in the user manual. Average recombination rates were computed using the stats program from the package, using option -burn 20 to discard the first 20 samples, corresponding to a burn-in of 20 *×* 5, 000 = 100, 000 iterations. The interval program being very slow to converge when the true recombination rate is zero, it was not run in these conditions for a sample size of 50 diploids. The corresponding estimate were recorded as missing data. The genome-average estimates were calculated using the average of all loci recombination rates, weighted by the distance between the corresponding SNPs.

#### 2.2.4 Pyrho

The *pyrho* software (version 0.1.6) was run using the recommended parameters [Spence and Song, 2019]. Two likelihood tables were generated, using constant or decreasing population sizes. In both cases, the true known demography was used. In the case of declining population sizes, 20 time-intervals of piece-wise constant population sizes were considered, starting 1,000 generations in the past. The population sizes *N_t_* of each interval *t* were computed using the formula

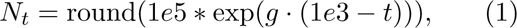

where the growth rate 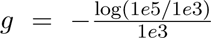 (exponential decline for a population of 100,000 diploids 1,000 generations ago, to a size of 1,000 diploids at present).

The impact of hyper-parameter choice was assessed using a grid of three block penalties and three window sizes. We found that the default parameters offered a good compromise across all criteria and kept them. The genome-average estimates were calculated using the average of all loci recombination rates, weighted by the distance between the corresponding SNPs.

#### 2.2.5 heRho

VCF files were converted to *heRho* input files using the provided heRho_tally_pairwise_counts_vcf.py script. *heRho* was then run using default parameters [Setter et al., 2022]. We reported two recombination rates from the *heRho* output: the crossing-over rate (kappa parameter) and the total recombination rate (kappa + omega), which includes GC events.

### 2.3 Statistical analysis

The estimated recombination rates were plotted against the true simulated crossing-over rate. All analyses were done in R (version 4.1.0) [R Core Team, 2021] with the *ggplot2* package (version 3.4.2) [Wickham, 2016].

## 3 Results

The comparison benchmark used in all the following consists of (i) simulating individual sequences according to a demographic scenario and recombination landscape, (ii) using the generated data to infer the average genome recombination rate (hereby noted *ρ*) using several population genomics methods, and (iii) comparing the resulting estimate to the true value used to simulated the data. The simulated sequences always consist of a unique chromosome of size 10 Mb and 10 replicates. The number of individuals used for inference, the demographic scenario, recombination landscape and inference method vary in each experiment.

### 3.1 Inference under an ideal scenario

We first consider an ideal scenario where the data is generated using a model very close to the inference model. The generative model consists of a standard coalescent with recombination, with constant population size in time and along the genome (no selection), and a flat recombination landscape (homogeneous recombination). We compared two SMC-based methods: the multiple sequentially coalescent (version 2), *MSMC* [Schiffels and Wang, 2020], and the integrative sequentially Markov coalescent, *iSMC* [Barroso et al., 2019]. For this comparison, *iSMC* was used with a single recombination class so that its model is identical to *MSMC*. As a comparison, two linkage disequilibrium (LD) based methods were also included: *LDhat* [McVean et al., 2004] and *Pyrho* [Spence and Song, 2019], a more recent re-implementation of the *LDhat* method. Lastly, we included the *heRho* method [Setter et al., 2022], which estimates recombination rates from the distance patterns of heterozygous sites.

Under this ideal scenario, most methods recover the recombination rate with high accuracy (Figure 1). LD-based methods slightly underestimate the recombination rate when the sample size is small (5 diploids) and the recombination rate is high. *heRho* accurately recovers the CO rate but tends to estimate a non-null rate of GC when the total recombination rate is low. Furthermore, *heRho* estimates have a larger variance when the sample size is small.

**Figure 1:**
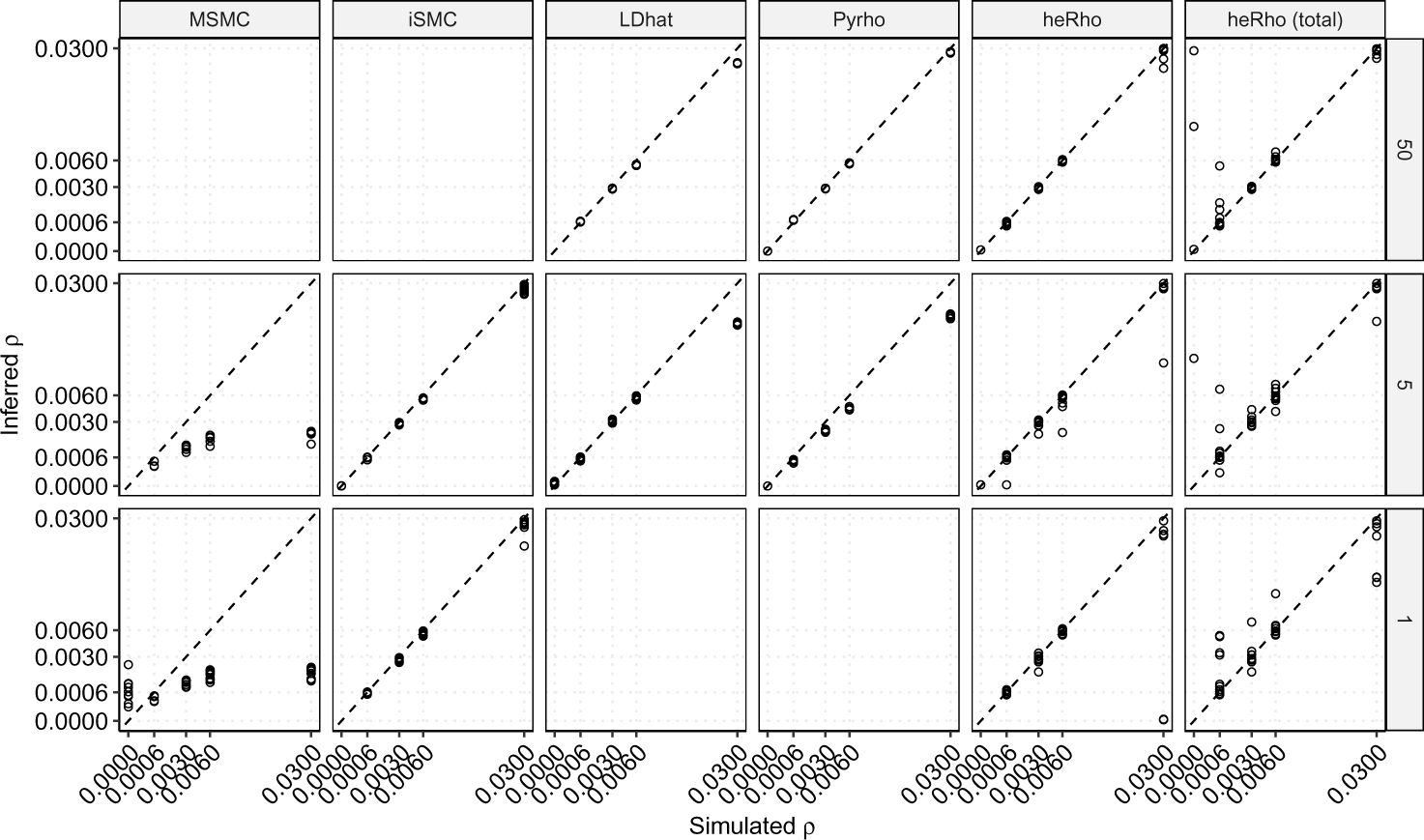
Inference of the genome-wide recombination rate under an ideal scenario of constant population size and flat recombination landscape. The inferred rate is plotted on the y-axis for distinct methods (column facets) and sample size (row facets, number of diploid individuals) as a function of the true rate on the x-axis. Both axes are plotted using a square-root scale, and the 1:1 diagonal is plotted as a dashed line. Estimates for the *iSMC* and *heRho* methods, sample size 1, and recombination rate 0 are above 0.03 and are not shown. SMC methods cannot be run on large sample sizes (such as *n* = 50, while LD-based methods cannot be run on single diploid genomes (*n* = 1).

*MSMC* estimates are very poor (overestimation of low / underestimation of high values). The difference between *MSMC* and *iSMC* is particularly surprising, given the strong similarity of the underlying model. Besides their independent implementations, the *iSMC* and *MSMC* methods differ in the demographic model specification (skyline model for *MSMC* vs splines for *iSMC*) and the optimization algorithm. The first point is not expected to play any role in the observed differences since the true population size is constant. As a control, we observe that *MSMC* recovers the true demography with fairly good accuracy with the exception of the largest recombination rates, although with large variance in the more recent and most distant past (Supplementary Figure 1). Several replicates fail to converge; *MSMC* is unable to converge in any of the 10 replicates when the true recombination rate is zero, and the proportion of failure decreases when the recombination rate augments (Supplementary Figure 1). Convergence issues arise when the analyzed genomes are too short, and following the *MSMC* authors’ recommendation, we reduced the number of time intervals in the skyline model to improve the performances [Schiffels and Wang, 2020]. Using half of the default time intervals (see sec:materials:methods:inference) provides a performance gain when the recombination rate is not zero, reducing the variance in the inferred population size and reducing the number of convergence failures. However, it has no impact on the recombination rate estimation (Supplementary Figure 2). Doubling the number of iterations in the Baum-Welch algorithm or starting from a different initial value does not lead to any improvement in the recombination estimation (Supplementary Figure 2).

SMC-based approaches are typically applied to a small number of genomes, possibly as little as a single diploid individual (or two haploid genomes). The estimation of *ρ* does not show any noticeable difference when a single unphased diploid (two haploid genomes) is used compared to five diploids (Figure 1). However, differences occur when the true rate is zero. *ρ* is erroneously found to be high when using a single individual, the overestimation being much stronger for *iSMC* than for *MSMC*. This is due to the sample sharing a unique ancestor for all positions, preventing the fit of a distribution of divergence times. When more individuals are used, *MSMC* fails to converge when the true rate is zero, while *iSMC* returns a very low value. We note that the key parameter here is the number of chromosomes rather than the actual number of individuals. The composite likelihood approach generates identical likelihood values whether two chromosomes come from the same or two distinct individuals since the Markov chain is reset and the likelihoods for each chromosome are multiplied. Even if the intra-chromosome recombination rate is zero, the inter-chromosomal rate would be equal to 0.5 and distinct chromosomes will have different ancestors, resulting in a better fit of the model. Finally, we note that the *ρ/θ* ratio, where *θ* = 4 *· Ne · u* = 0.005, varies from 0.12 to 6 throughout the simulations, showing that SMC-based approaches can infer the genome-average recombination rate even in situations where *ρ/θ >* 1 and some recombination events leave no trace in the data.

In conclusion, we note that under an ideal scenario:

1. The SMC permits the inference of the genome-average recombination rate with good accuracy and with as little as one diploid individual.
2. When the true rate is high and the sample size is small, the estimates are less biased than those of the LD-based methods.
3. For low recombination rates, the estimation variance is lower for SMC-based estimates than for the heterozygosity-based method.
4. A standard maximum likelihood optimization procedure should be used, as the Baum-Welch expectation-maximisation algorithm does not permit obtaining accurate estimates.
5. If the data set consists of a single chromosome of a single individual and the true recombination rate is zero, SMC methods will provide erroneous estimates.

### 3.2 Inference under a heterogeneous recombination landscape

We use the same setup as in the ideal scenario, but now the recombination landscape is no longer homogeneous along the chromosome. Segments of uniform recombination rates are randomly generated, their lengths are taken from a geometric distribution and their values are exponentially distributed (see sec:materials:methods:simulations). In addition to the methods tested in the previous case, we add the *iSMC* method with five recombination classes (noted *rhoSMC* in the following, Barroso et al. [2019]). The *rhoSMC* model includes the genome average recombination rate as a parameter that can be estimated by maximum likelihood (referred to as maximum likelihood estimate, MLE, in the following), similar to the *iSMC* simpler model. In addition, *rhoSMC* allows computing a posterior average estimate (referred to as posterior estimate in the following, see sec:materials:methods:inference). Under this scenario, all methods assuming a homogeneous recombination rate underestimate the genome average rate (*MSMC*, *iSMC*, and *heRho*), while LD-based methods, which infer a recombination map, show the same performance as when the landscape is flat (only slightly under-estimating high recombination rates when the sample size is small). While *heRho* generally underestimates the CO rate, in this scenario we note that its total recombination rate estimate is closer to the true rate, suggesting that *heRho* interprets some of the signatures of variable recombination rate as GC events. The *rhoSMC* method reports unbiased estimates (Figure 2). The MLE and posterior estimates were similar, the posterior estimate being slightly superior when the recombination rate is low. We note that all methods with the exception of *heRho* achieve similar performance when simulating under a population size ten times smaller, resulting in a ten times lower genetic diversity (Supplementary Figure 3). Conversely, *heRho* estimates have a much larger variance under these conditions.

**Figure 2:**
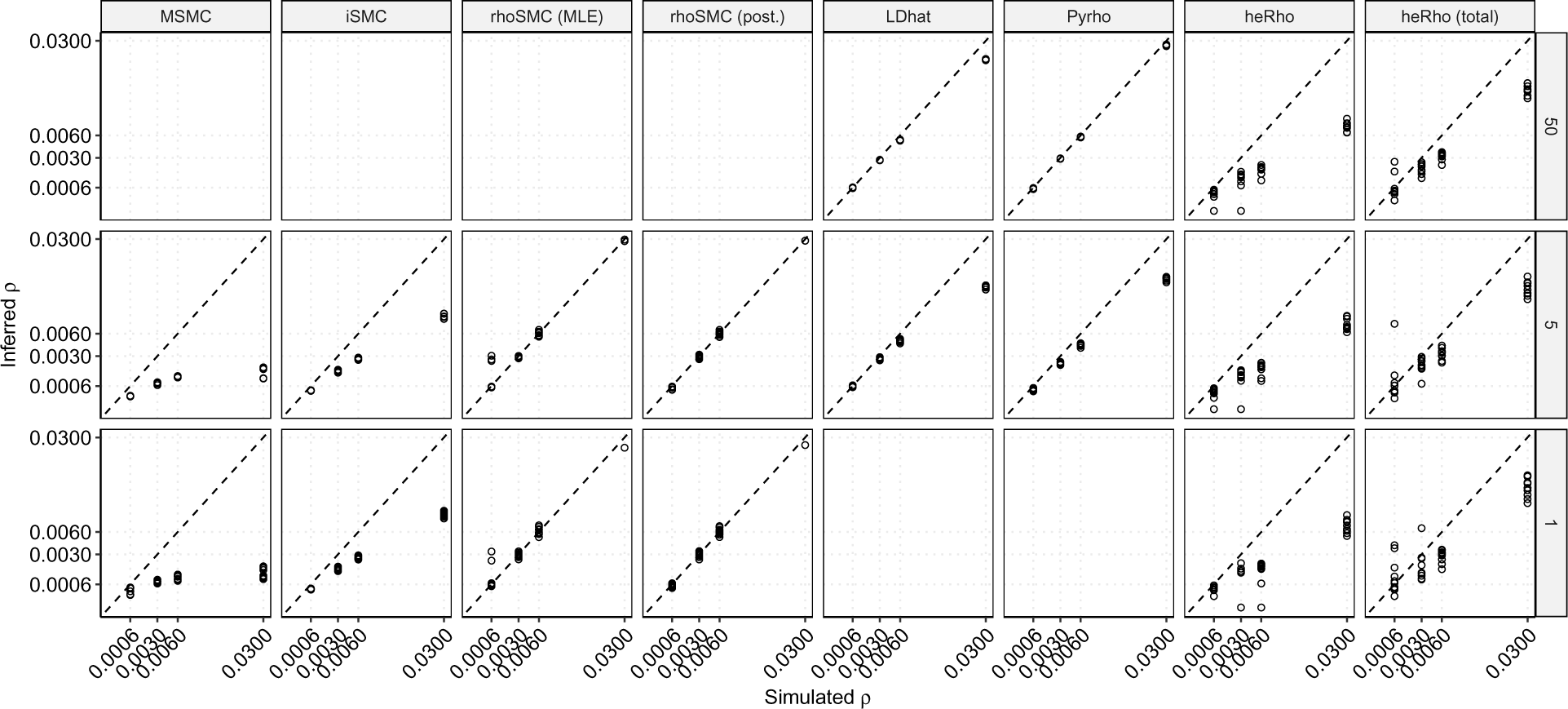
Inference of the genome-wide recombination rate under constant population size and a heterogeneous recombination landscape. Legend as in Figure 1.

The *rhoSMC* method assumes a Markov process of recombination rate category change along the genome. To assess its performance when the recombination landscape does not follow this auto-correlation pattern, the analysis was repeated on a recombination landscape taken from the deCODE human recombination map (see sec:materials:methods); the results and conclusions proving to be qualitatively similar. We note a higher variance between replicates for the inference with *rhoSMC* and a larger underestimation with *heRho* on this landscape, compared to the randomly generated one (Supplementary Figure 4).

Next, we investigated the performance of *rhoSMC* – which assumes a heterogeneous recombination landscape – when the true landscape is homogeneous. The maximum likelihood estimator appeared to be biased under this scenario, while the posterior average estimate recovered the true recombination rate, providing the rate is not zero (Supplementary Figure 5). In this case, as shown before, the estimators are biased if only a single chromosome is used. This result suggests that some model testing should be done before interpreting the estimated values.

The heterogeneous *rhoSMC* model (M1) can be compared to the homogeneous *iSMC* model (M0) by setting the number of recombination categories to 1. The presence of recombination heterogeneity can be assessed by comparing the two models, either by means of Akaike’s information criterion (AIC) or using a likelihood ratio test (LRT). When computed on a homogeneous data set, the AIC difference between the two models is distributed between −10 and 10, independently of the recombination rate (Supplementary Figure 6A), although it reaches -1e6 when the recombination rate is 0 and 5 individuals are used. In comparison, the AIC differences reach −1000 when applied to a heterogeneous landscape. While AIC seems able to disentangle the two models, the threshold to use is ambiguous, meaning that simulations tailored for the data set to analyse are probably required to make a proper goodness-of-fit test. Similar conclusions are reached when considering the LRT, where lower P-values are obtained on heterogeneous data sets, but P-values lower than 1%, and even 0.01%, are observed with homogeneous data sets (Supplementary Figure 6B). Lastly, the presence of GC mimics rate heterogeneity, leading to a stronger signal for rejecting M0 when the true landscape is homogeneous (Supplementary Figure 6C and D). We further discuss the effect of GC in sec:results:gc.

We conclude that under a heterogeneous recombination landscape:

1. Assuming a homogeneous recombination landscape in the SMC leads to an under-estimation of the average rate, even when a pure maximum likelihood optimization framework is used.
2. Accounting for the recombination rate variation in the SMC permits an unbiased inference of the average rate, even when the underlying recombination landscape differs from the underlying model of rate variation.
3. Maximum likelihood estimates of the average recombination rate are biased when a heterogeneous model is used, and the true landscape is homogeneous. Posterior average estimates show better performance under these conditions unless the recombination rate is zero.
4. The heterozygosity-based inference strongly underestimates the average recombination rate when the landscape is heterogeneous.

### 3.3 Inference under non-constant demography

The population recombination rate depends on the product of the CO rate and the population size, which may vary over time. Assuming that the average recombination rate is constant over the period since the most recent common ancestor of the sample is a relatively reasonable assumption. Conversely, variation in population sizes over this time scale may have a strong impact on the rate estimate [Kamm et al., 2016]. We simulated data under the variable recombination scenario and a non-constant demography scenario. The population size was set to 100,000, as before, but it exponentially decreased starting 1,000 generations ago to reach a population size of 1,000 at present. Under this decreasing population size scenario, methods which assume a flat demography (*LDhat*, *heRho*) systematically under-estimate the genome-average population recombination rate (Figure 3. The LD-based method *Pyrho* provides accurate estimates since the underlying demography is taken into account when computing the likelihood table with the *LDpop* method [Kamm et al., 2016]. The *rhoSMC* method also recovers the true recombination rate as it jointly estimates demography with the recombination landscape.

**Figure 3:**
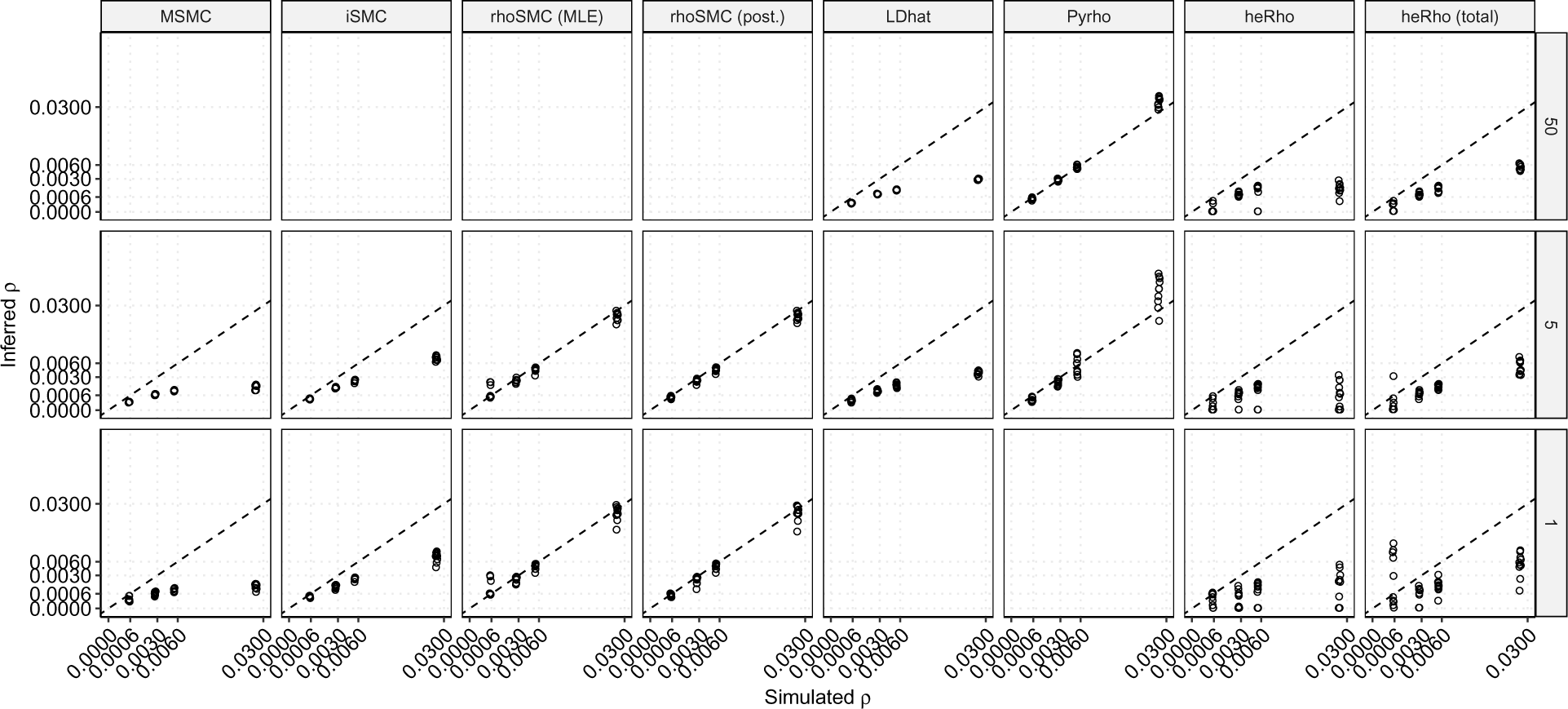
Inference of the genome-wide recombination rate when population sizes decline. Simulations under a variable recombination rate, legends as in Figure 2.

**Figure 4:**
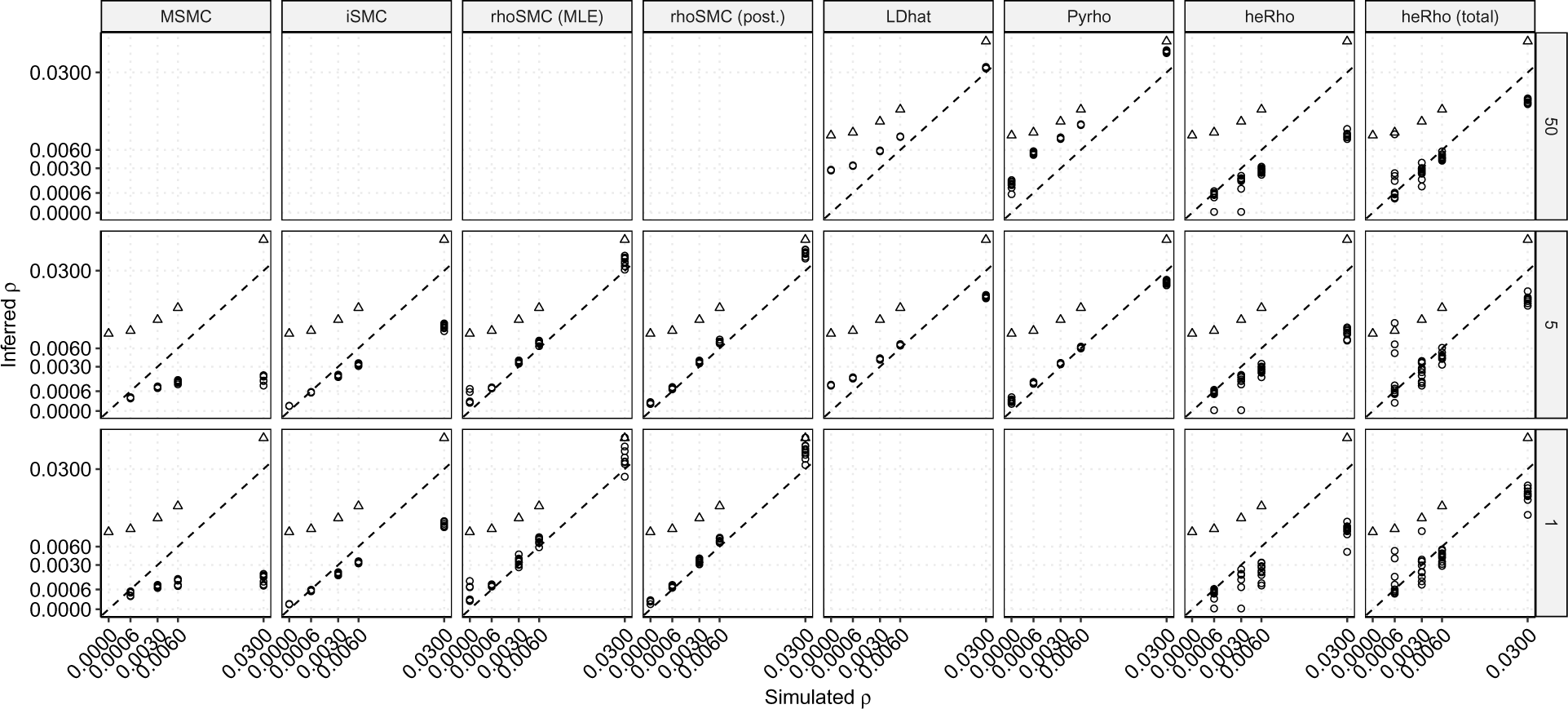
Inference of the genome-wide recombination rate in the presence of recombination hotspots. Simulations under a constant population size. The x-axis shows the background rate. Circles show the inferred *ρ*. Triangles show the true genome-average recombination rate as a weighted average of background and hotspots. Other legends as in Figure 1.

We observed consistent results under scenarios where the population size increased (Supplementary Figure 7. In the first scenario, we considered an exponential increase 1,000 generations ago from a size of 1,000 individuals to a size of 100,000 at present. In the second scenario, an increase from 50,000 to 200,000 is considered. The resulting genetic diversity is reduced to 15% in the first scenario and 50% in the second, compared to a scenario with a constant population size of 100,000. Under these scenarios, *LDhat* overestimates the population recombination rate while *Pyrho* and *iSMC* properly account for the underlying demography. *Herho*’s performance is not significantly affected in the second scenario, but the estimates’ variance becomes extremely large in the first scenario because of the reduced diversity (see also Supplementary Figure 3). The estimates of the total recombination rate in particular become unreliable.

Conclusions:

**1.** Ignoring population size variation leads to biased estimates of the recombination rate.
2. Accounting for demography when computing the likelihood table permits accurate inferences using LD-based methods.
3. The *rhoSMC* method accurately infers the genome-average recombination rate when the demography is not constant.

### 5.1 Inference in the presence of recombination hotspots

In many species, the recombination landscape is shaped by recombination hotspots, short regions with very high recombination rates. Their localized nature and the high number of recombination events make them difficult to infer and require a high-resolution recombination map. To assess the accuracy of the inference of the genome-average recombination rate in the presence of hotspots, we simulated random regions of 0.5 to 2 kb with a recombination rate of *ρ* = 0.35 (on average 70x the background rate) spaced by uniform regions of 50 kb on average (see Methods). We observed that all methods underestimate the average recombination rate under this scenario (Supplementary Figure 4). Only LD-based methods capture a partial signal from the recombination hotspots when a large sample size is available. Methods applied to small sample sizes (SMC and LD-based methods) only infer the background recombination rate. Given their pairwise nature, SMC-based methods are in particular blind to the presence of recombination hotspots.

### Conclusions

1. Only LD-based methods and large sample sizes capture the effect of recombination hotspots.
2. SMC-based methods efficiently measure the background recombination rate.

### 5.2 Inference in the presence of gene conversion

We finally assessed the impact of GC events on the inference of the genome-average recombination rate. The simulation setup is kept identical to the above, but a variable proportion of GC events is now implemented. We kept the average total recombination rate (CO + GC events) equal to *ρ* = 0.06 and implemented proportions of GC of 0%, 10%, 50%, 90% and 100%, with a constant track length of 300 bp.

Most of the tested methods performed poorly when GC is present (Figure 5). *heRho* is the only method able to properly infer the total and CO recombination rate. We note, however, that this is only the case when the recombination landscape is flat, and the population size is constant, the recombination rate being underestimated otherwise. LD-based methods tend to infer an intermediate rate, lower than the total recombination rate but higher than the CO rate. Under a flat recombination landscape, the *iSMC* method also infers an intermediate rate, providing GC events do not represent 100% of the recombination events. The *rhoSMC* method proved to be highly sensitive to the presence of GC events, resulting in an overestimation of the recombination rate. This indicates that the signature of GC events is similar to that of multiple CO events. We note that the estimated *ρ* is close to the predicted rate when GC events would be counted as two CO events (+1 slope on figure 5).

**Figure 5:**
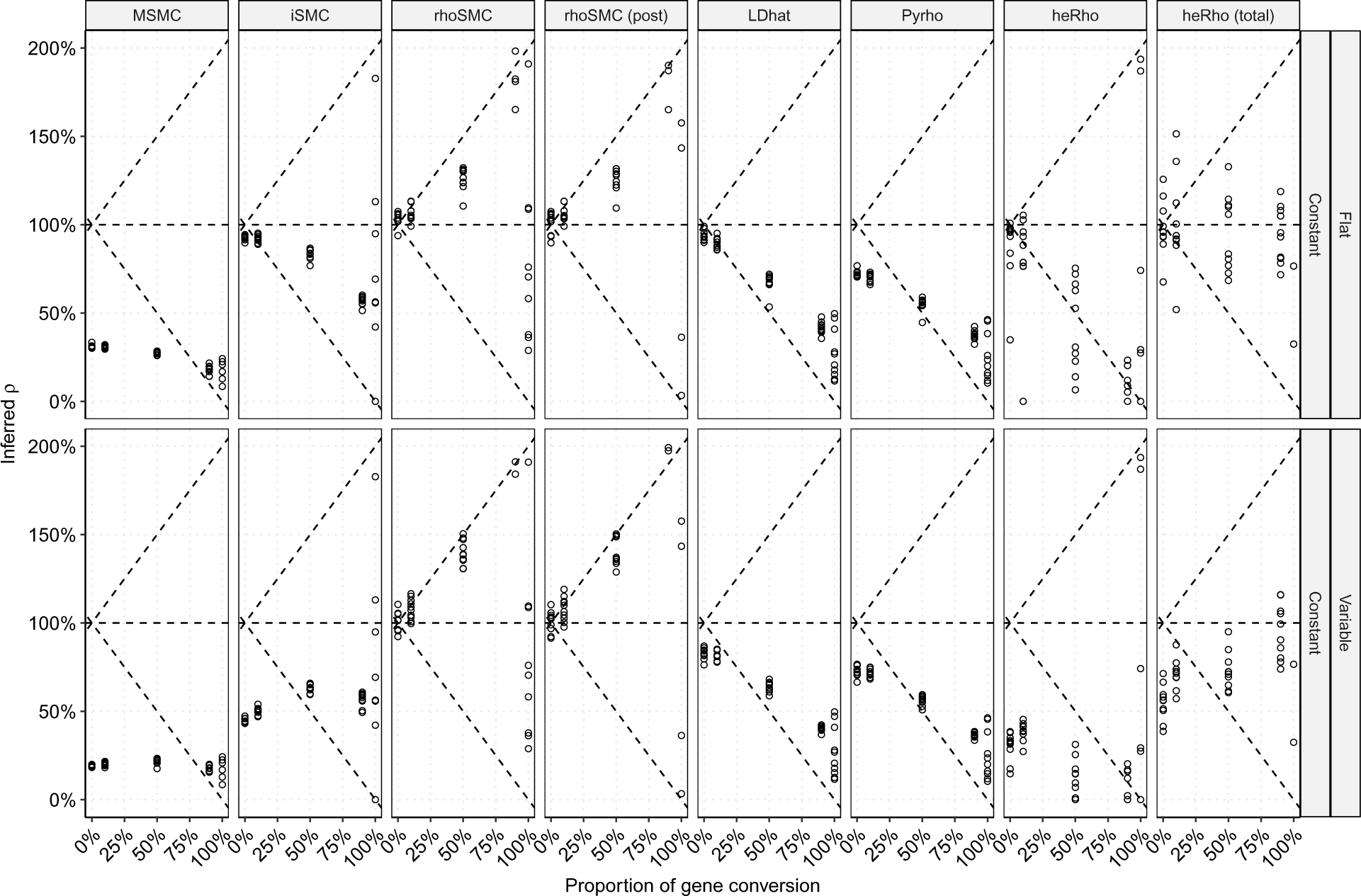
Inference of the genome-wide recombination rate in the presence of gene conversion (GC). Simulations under a total average recombination rate of *ρ* = 0.06 with various proportions of GC events (x-axis). The y-axis reports the ratio of the inferred recombination rate to the true one as percentages. Facet rows display various scenarios. Sample size is equal to 5 diploids in all scenarios. Results for sample sizes of 1 are shown in Supplementary Figure 8 and 50 in Supplementary Figure 9. Horizontal dash lines correspond to the true relative recombination rate (100%) and the −1 slope dashed line shows the CO rate as a proportion of the total recombination rate. The +1 slope dashed lines show the theoretical total recombination rate when GC events count as two recombination events.

### Conclusions

1. Gene conversion has a strong impact on the inference of the genome-average recombination rate.
2. The signal of gene conversion is intermingled with that of recombination rate variation.
3. The *heRho* method can disentangle gene conversion and crossing-over events when the recombination rate is homogeneous but is biased when it is heterogeneous or when the population size is not constant in time.
4. *rhoSMC* properly captures the heterogeneity of the recombination landscape but overestimates the recombination rate in the presence of gene conversion.

## 6 Discussion

While estimating the genome average recombination rate *ρ* = 4 *· Ne · r*, a single parameter, may seem a simple endeavour in comparison to inferring a detailed recombination map, the task is challenging because *r* varies spatially and *Ne* temporally. (Not to mention the possibility that *r* may also vary temporally and *Ne* spatially, two possibilities that we have not investigated in this study.) LD-based methods (in particular the most recent ones such as *Pyrho*) that account for non-constant demography proved to provide the most accurate estimations under all tested scenarios. However, they require relatively large sample sizes. Conversely, SMC-based methods offer a powerful alternative when as little as one unphased diploid genome is available. Such methods properly account for the demography of the underlying population, which is jointly estimated with recombination rate parameters. Two factors appear important when inferring recombination rates with SMC models: using a full maximum-likelihood estimation procedure instead of the Baum-Welch algorithm and properly accounting for the heterogeneity of the recombination rate.

SMC models, however, are not able to capture the signal of recombination hotspots and only infer the background recombination rate. As recombination hotspots represent by nature very fine-scale variation, their signal may only be captured with large sample sizes. As of today, only LD-based methods are powerful enough to capture this signal.

We further showed that the presence of GC may significantly bias the recombination rate inference. LD-based methods essentially capture the CO rate and some of the GC events. SMC-based methods, however, show a strong bias as they tend to infer two CO events for one GC event, leading to a significant overestimation of the recombination rate in case the proportion of GC events is high. This result is particularly concerning as the relative rate of GC vs. CO has been suggested to vary extensively between genomic regions and could be as high as 100 [Padhukasahasram and Rannala, 2013]. The *heRho* method is currently the only one that can accurately infer both the CO and total recombination rate in the presence of GC. However, in its current implementation, it can only do so when the recombination landscape is homogeneous and the demography constant – a largely unrealistic scenario. In the original publication [Setter et al., 2022], the authors assess the impact of rate heterogeneity by mixing two homogeneous datasets with distinct recombination rates and did not find a significant impact on the estimation. Here, we used a more realistic scenario where the recombination rate takes values from an exponential distribution or from an empirical distribution (the deCODE dataset) and found a significant impact of recombination rate heterogeneity on the inference of the average rate. While non-constant demography may be incorporated in the modelling, the signal of GC and variable recombination rate may be difficult to disentangle. In that respect, a convergence between the correlation of heterozygosity and SMC-based methods, possibly by incorporating GC into the SMC model framework, appears as a promising avenue.

## 7 Data availability

All the necessary code to reproduce the analyses and figures in this study, as well as intermediate result tables, are available at https://gitlab.gwdg.de/molsysevol/smc-benchmark. The version used in this manuscript is also available at FigShare, DOI: 10.6084/m9.figshare.24069270. Supplementary File S1 provides a notebook with the R code to generate all figures.

## Acknowledgments

The author would like to thank Gustavo Barroso for discussions on this project. This project emerged from discussions with Gwenael Piganeau and Victor Loegler. The idea of assessing the impact of gene conversion on recombination inference was suggested by Richard Durbin, after a presentation by the author at the Pop-Group meeting in 2022.

## 10 Supplementary Material

**Supplementary Figure 1:**
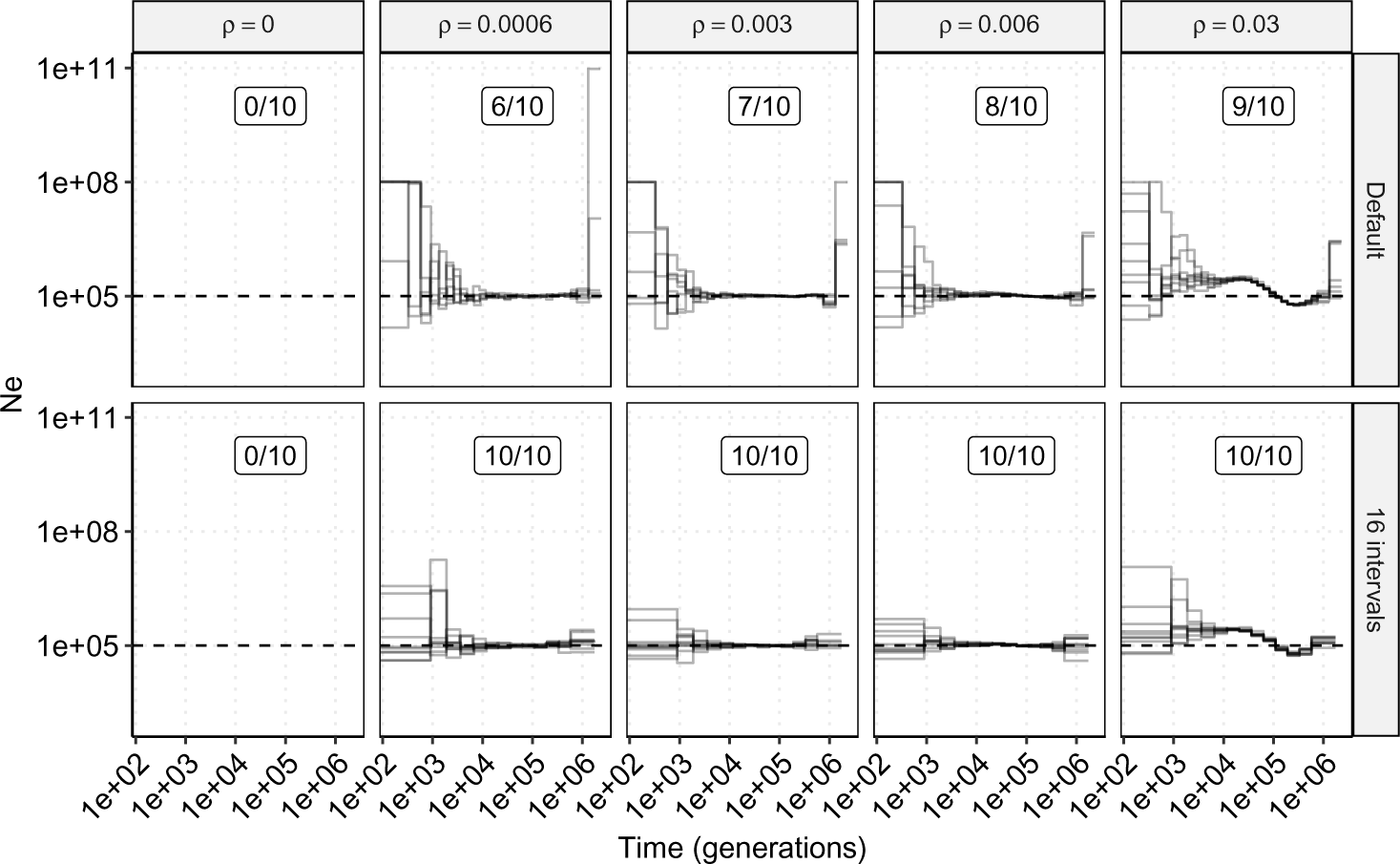
Demography inference using MSMC. Ten datasets of five diploid individuals were simulated for each parameter-method combination under a constant population size scenario (horizontal dashed line *Ne* = 100, 000). Column facets depict distinct recombination rates and row facets compare the inference method: default parameters (top) or reduced time intervals (bottom). Framed numbers show the proportion (out of 10) of replicates where the optimization converged.

**Supplementary Figure 2:**
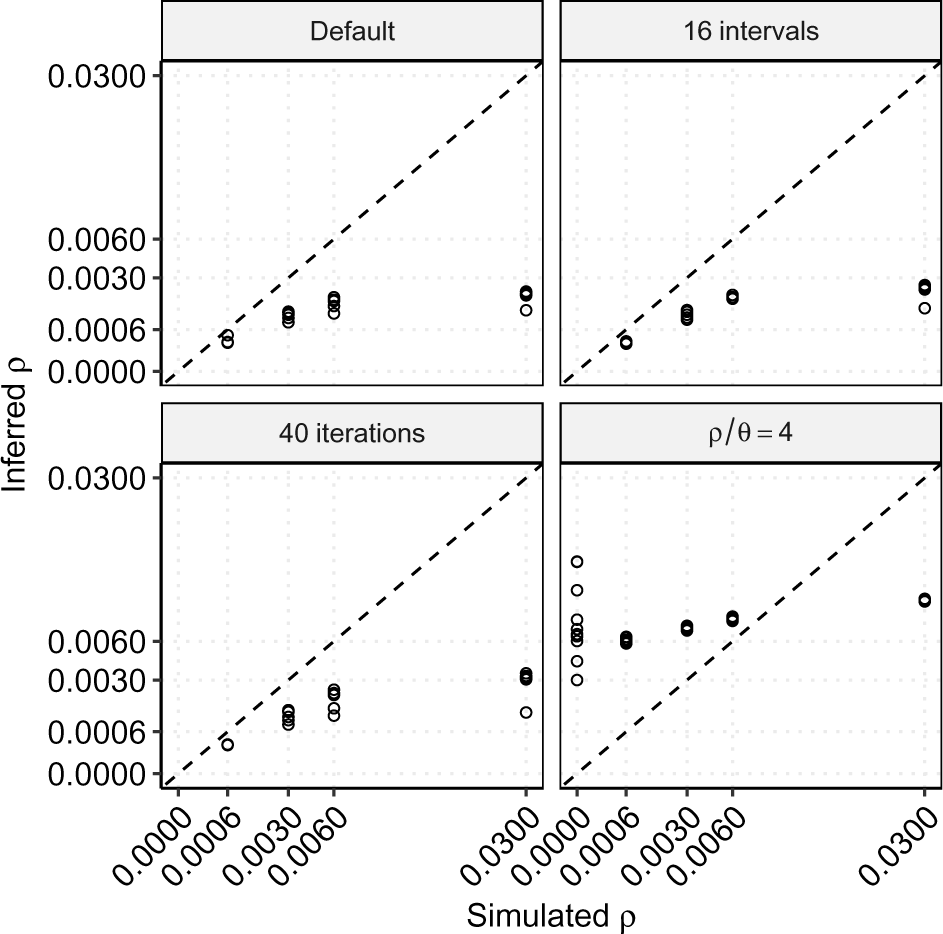
Impact of the MSMC model and optimization procedure on the inference of the genome-average recombination rate. Five diploid individuals were used in each simulated dataset. Legend as in Figure 1.

**Supplementary Figure 3:**
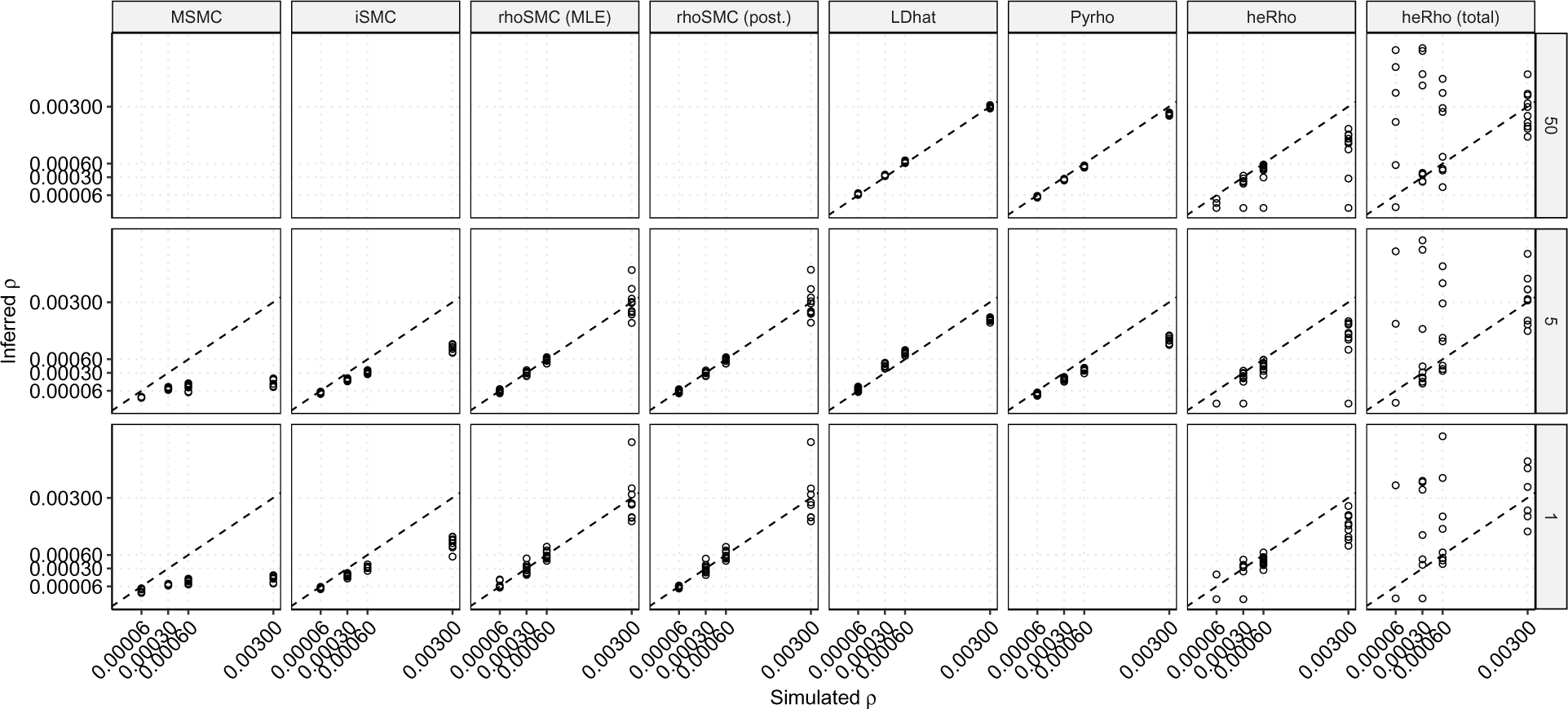
Inference of the genome-wide recombination rate under a heterogeneous recombination landscape and constant population size equal to 10,000 individuals. Legend as in Figure 2, which used a population size of 100,000 individuals.

**Supplementary Figure 4:**
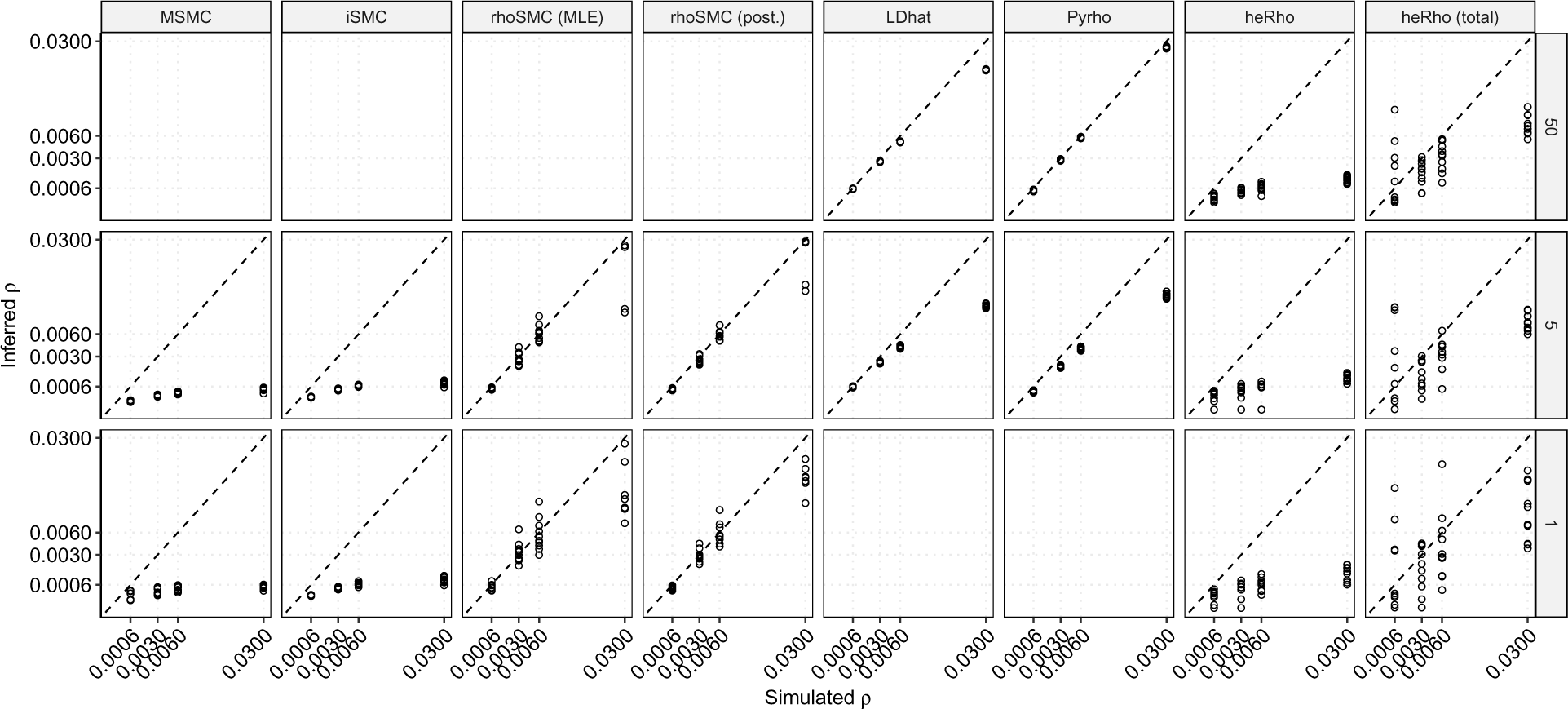
Inference of the genome-wide recombination rate under constant population size and a heterogeneous recombination landscape, taken from the DECODE recombination map. Legend as in Figure 2.

**Supplementary Figure 5:**
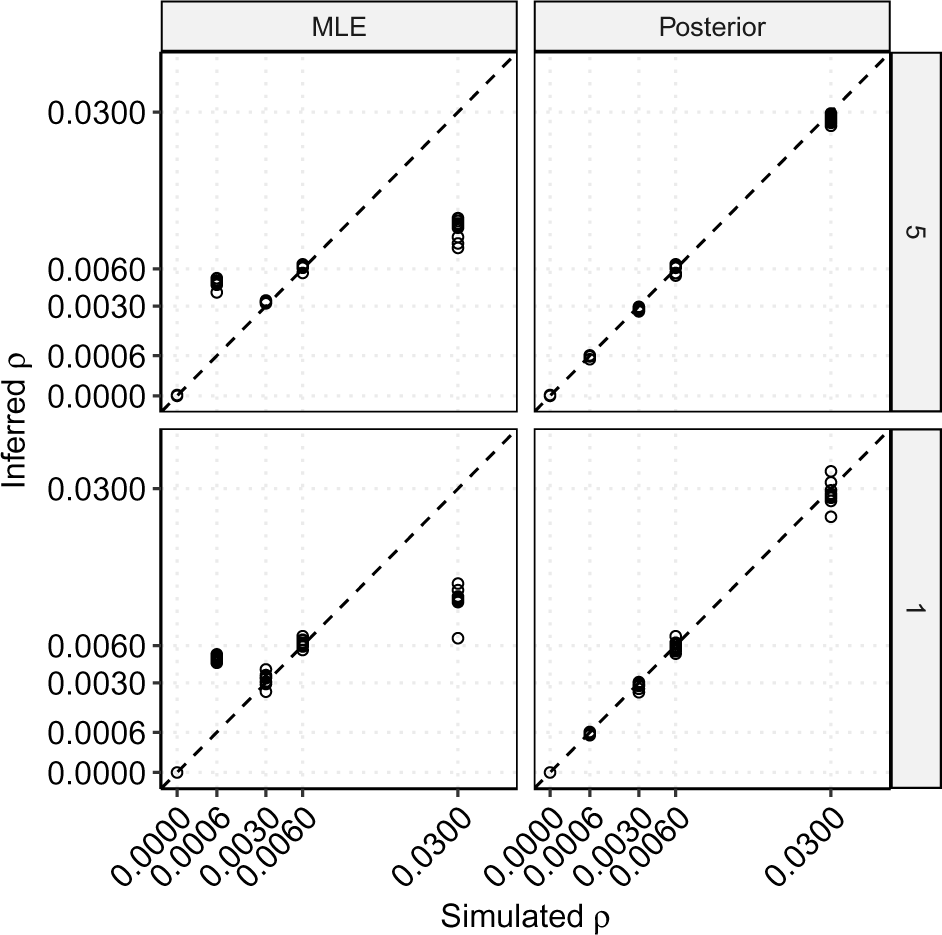
Performance of the *rhoSMC* method when the true recombination landscape is homogeneous. MLE: maximum likelihood estimates. Posterior: posterior average estimates. Facet rows show the sample size (number of diploid individuals). When a single individual is used, and the true recombination rate is 0, *rhoSMC* erroneously returns high estimates above 0.04 in 6 replicates out of 10, which are omitted in this figure.

**Supplementary Figure 6:**
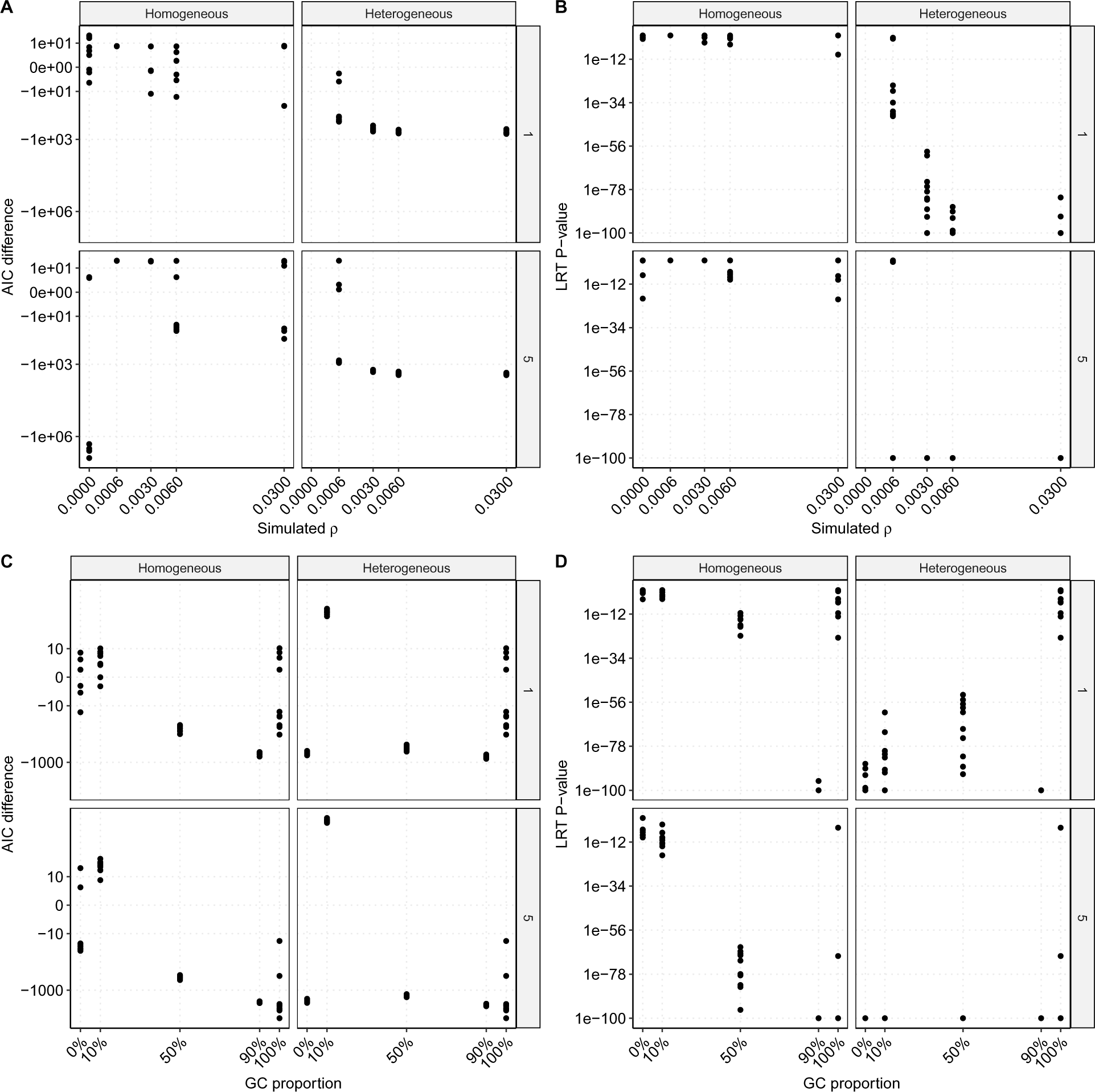
Model comparisons on homogeneous and non-homogeneous recombination landscapes, with and without gene conversion. Column facets: simulations under a homogeneous (flat) or heterogeneous (variable) recombination landscape. Row facets: number of diploid individuals used for inference. A,B: impact of the recombination rate. C,D: impact of the proportion of gene conversion (GC). A,C: Akaike’s information criterion (AIC). B,D: likelihood ratio test (LRT).

**Supplementary Figure 7:**
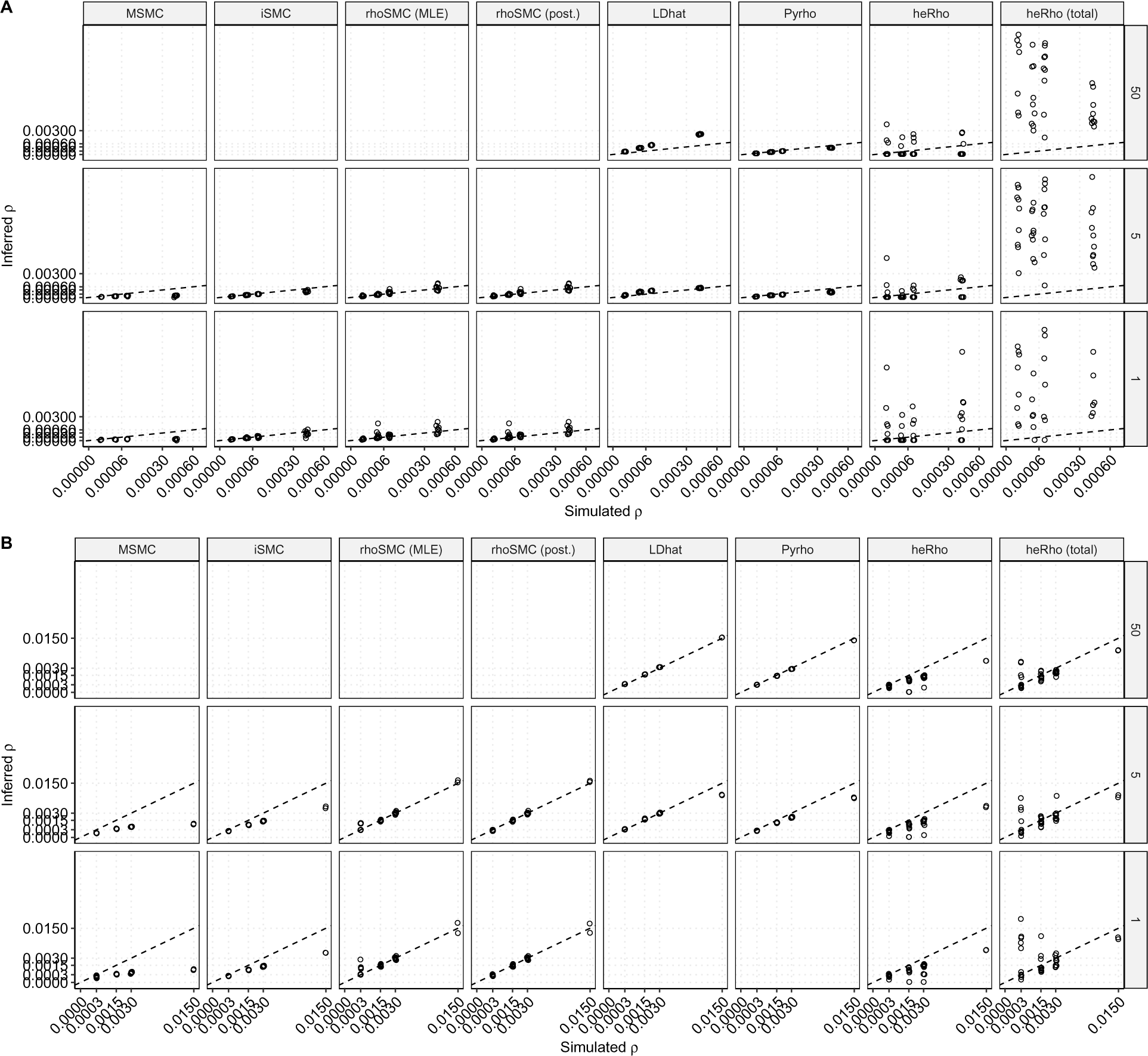
Inference of the genome-wide recombination rate when population sizes increase. A) Increase from a size of 1,000, 1,000 generations ago, to 100,000. B) Increase from a size of 50,000 to 200,000. Simulations under a variable recombination rate, legends as in Figure 3

**Supplementary Figure 8:**
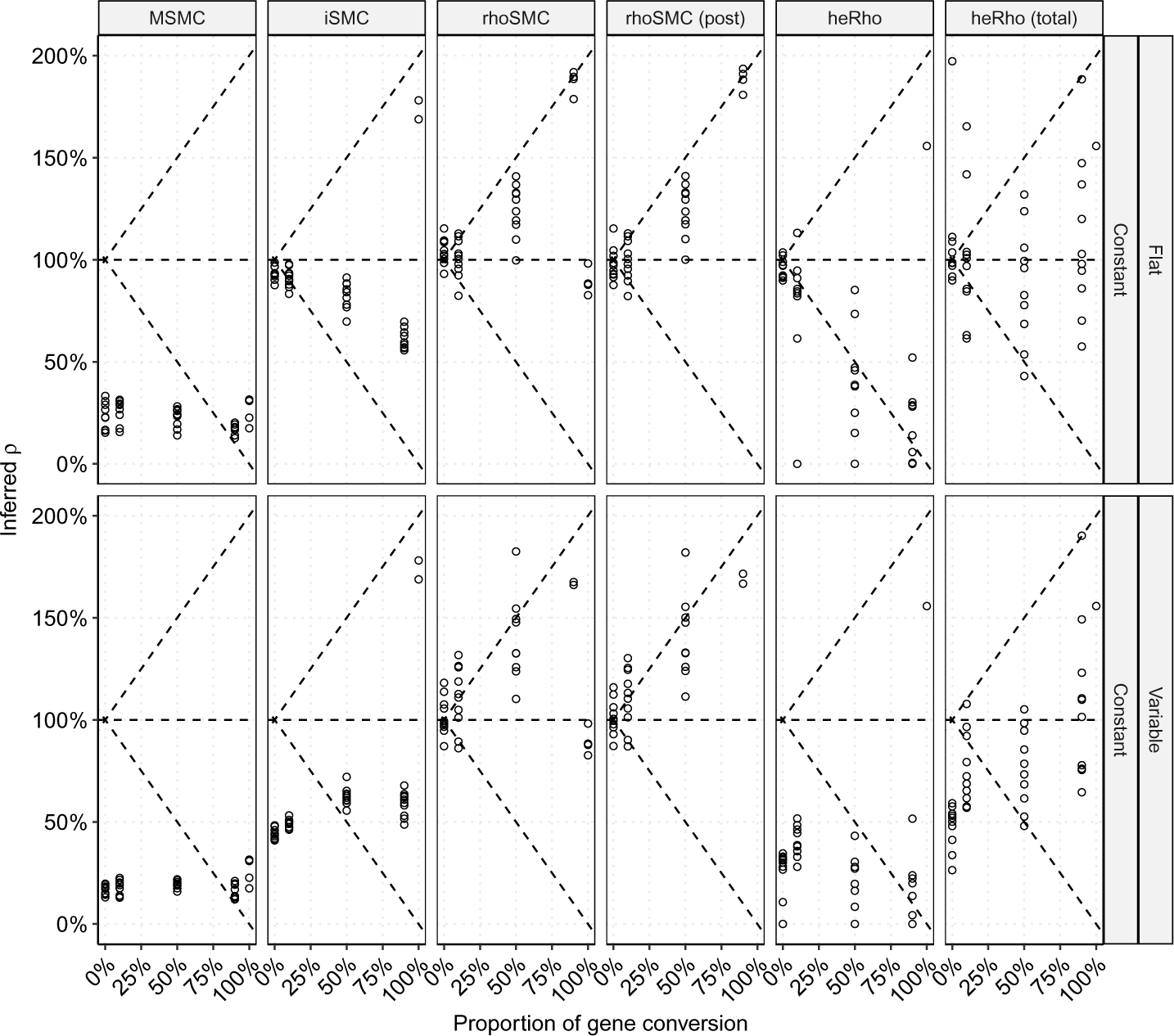
Inference of the genome-wide recombination rate in the presence of gene conversion. Inference from a single diploid individual. Legend as in Figure 5

**Supplementary Figure 9:**
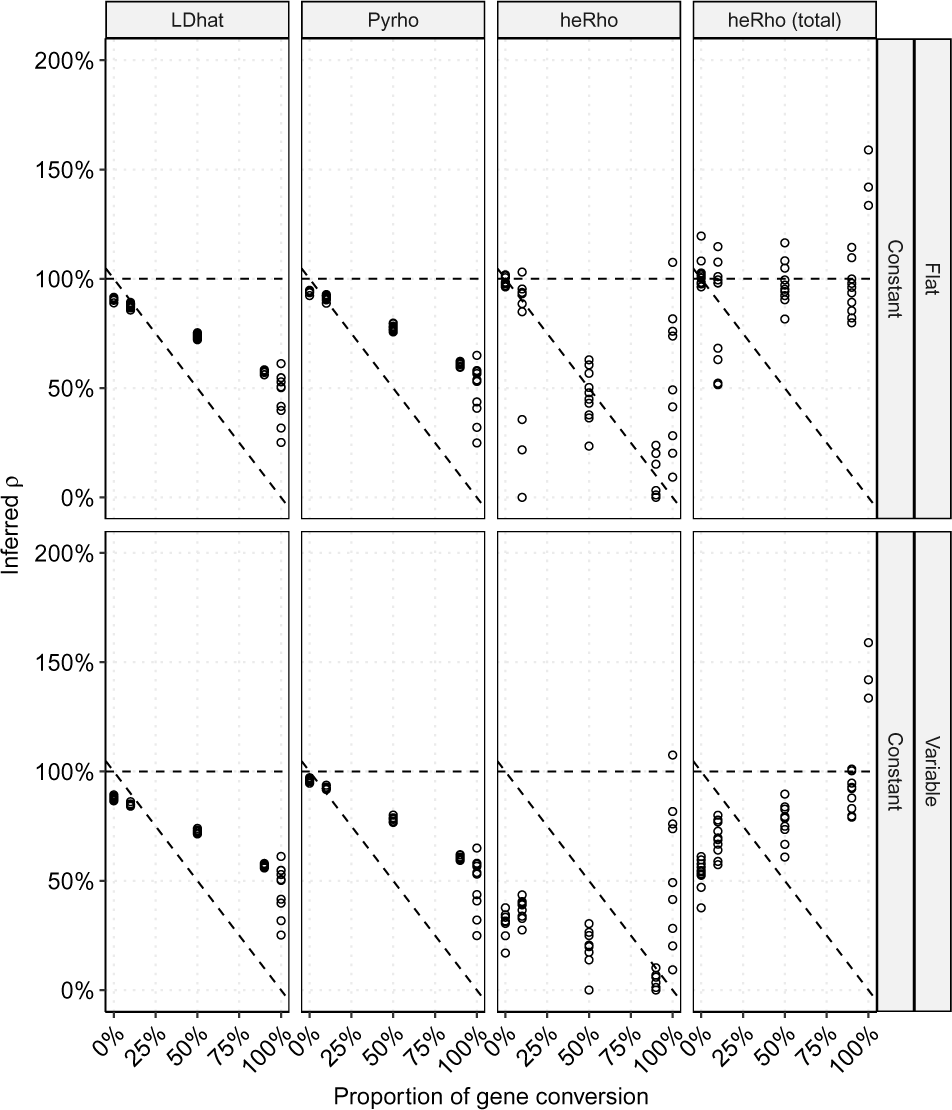
Inference of the genome-wide recombination rate in the presence of gene conversion. Inference from 50 diploid individuals. Legend as in Figure 5

## References

G. V. Barroso, N. Puzovíc, and J. Y. Dutheil. Inference of recombination maps from a single pair of genomes and its application to ancient samples. PLOS Genetics, 15(11): e1008449, Nov. 2019. ISSN 1553-7404. doi: 10.1371/journal.pgen.1008449.

F. Baumdicker, G. Bisschop, D. Goldstein, Gower, A. P. Ragsdale, G. Tsambos, S. Zhu, B. Eldon, E. C. Ellerman, J. G. Galloway, A. L. Gladstein, G. Gorjanc, B. Guo, B. Jeffery, W. W. Kretzschumar, K. Lohse, M. Matschiner, D. Nelson, N. S. Pope, C. D. Quinto-Cortés, M. F. Rodrigues, K. Saunack, T. Sellinger, K. Thornton, H. van Kemenade, A. W. Wohns, Y. Wong, S. Gravel, A. D. Kern, J. Koskela, P. L. Ralph, and J. Kelleher. Efficient ancestry and mutation simulation with msprime 1.0. Genetics, 220(3):iyab229, Mar. 2022. ISSN 1943-2631. doi: 10.1093/genetics/iyab229.

P. Danecek, A. Auton, G. Abecasis, C. A. Albers, E. Banks, M. A. DePristo, R. E. Handsaker, G. Lunter, G. T. Marth, S. T. Sherry, G. McVean, R. Durbin, and 1000 Genomes Project Analysis Group. The variant call format and VCFtools. Bioinformatics, 27(15): 2156–2158, Aug. 2011. ISSN 1367-4811. doi: 10.1093/bioinformatics/btr330.

J. Y. Dutheil. Towards more realistic models of genomes in populations: The Markov-modulated sequentially Markov coalescent. In E. Baake and A. Wakolbinger, editors, Probabilistic Structures in Evolution, pages 383–408. EMS Press, May 2021. ISBN 978-3-9854700-5-1. doi: 10.4171/ECR/17.

R. Epstein, N. Sajai, M. Zelkowski, A. Zhou, K. R. Robbins, and W. P. Pawlowski. Exploring impact of recombination landscapes on breeding outcomes. Proceedings of the National Academy of Sciences, 120(14):e2205785119, Apr. 2023. doi: 10.1073/pnas.2205785119. Publisher: Proceedings of the National Academy of Sciences.

B. Haubold, P. Pfaffelhuber, and M. Lynch. mlRho a program for estimating the population mutation and recombination rates from shotgun-sequenced diploid genomes. Mol Ecol, 19 Suppl 1(Suppl 1):277–284, Mar. 2010. ISSN 1365-294X. doi: 10.1111/j.1365-294X.2009.04482.x.

J. A. Kamm, J. P. Spence, J. Chan, and Y. S. Song. Two-Locus Likelihoods Under Variable Population Size and Fine-Scale Recombination Rate Estimation. Genetics, 203(3):1381– 1399, 2016. ISSN 1943-2631. doi: 10.1534/genetics.115.184820.

A. Kong, D. F. Gudbjartsson, J. Sainz, G. M. Jonsdottir, S. A. Gudjonsson, B. Richardsson, S. Sigurdardottir, J. Barnard, B. Hallbeck, G. Masson, A. Shlien, S. T. Palsson, M. L. Frigge, T. E. Thorgeirsson, J. R. Gulcher, and K. Stefansson. A high-resolution recombination map of the human genome. Nat. Genet., 31(3):241–247, July 2002. ISSN 1061-4036. doi: 10.1038/ng917.

H. Li and R. Durbin. Inference of human population history from individual whole-genome sequences. Nature, 475(7357):493–496, July 2011. ISSN 1476-4687. doi: 10.1038/nature10231.

G. A. T. McVean, S. R. Myers, S. Hunt, P. Deloukas, D. R. Bentley, and P. Donnelly. The fine-scale structure of recombination rate variation in the human genome. Science, 304 (5670):581–584, Apr. 2004. ISSN 1095-9203. doi: 10.1126/science.1092500.

B. Padhukasahasram and B. Rannala. Meiotic gene-conversion rate and tract length variation in the human genome. Eur J Hum Genet, pages 1–8, Feb. 2013. ISSN 1476-5438. doi: 10.1038/ejhg.2013.30. Publisher: Nature Publishing Group.

J. V. Peñalba and J. B. W. Wolf. From molecules to populations: appreciating and estimating recombination rate variation. Nat Rev Genet, 21(8):476–492, Aug. 2020. ISSN 1471-0064. doi: 10.1038/s41576-020-0240-1. Number: 8 Publisher: Nature Publishing Group.

R Core Team. R: A Language and Environment for Statistical Computing. R Foundation for Statistical Computing, Vienna, Austria, 2021. URL https://www.R-project.org/.

K. Rengefors, A. Kremp, T. B. Reusch, and M. Wood. Genetic diversity and evolution in eukaryotic phytoplankton: revelations from population genetic studies. Journal of Plankton Research, 39(2):165–179, Mar. 2017. ISSN 0142-7873. doi: 10.1093/plankt/fbw098.

S. Schiffels and K. Wang. MSMC and MSMC2: The Multiple Sequentially Markovian Coalescent. Methods Mol Biol, 2090:147–166, 2020. ISSN 1940-6029. doi: 10.1007/978-1-0716-0199-07.

D. Setter, S. Ebdon, B. Jackson, and K. Lohse. Estimating the rates of crossover and gene conversion from individual genomes. Genetics, 222 (1):iyac100, Sept. 2022. ISSN 1943-2631. doi: 10.1093/genetics/iyac100.

J. P. Spence and Y. S. Song. Inference and analysis of population-specific fine-scale recombination maps across 26 diverse human populations. Science Advances, 5(10):eaaw9206, Oct. 2019. doi: 10.1126/sciadv.aaw9206.

J. P. Spence, M. Steinrücken, J. Terhorst, and Y. S. Song. Inference of population history using coalescent HMMs: review and outlook. Curr. Opin. Genet. Dev., 53:70–76, Dec. 2018. ISSN 1879-0380. doi: 10.1016/j.gde.2018.07.002.

J. Stapley, P. G. D. Feulner, S. E. Johnston, A. W. Santure, and C. M. Smadja. Variation in recombination frequency and distribution across eukaryotes: patterns and processes. *Philos. Trans. R. Soc. Lond., B*, Biol. Sci., 372(1736): 20160455, Dec. 2017. ISSN 1471-2970. doi: 10.1098/rstb.2016.0455.

K. Theissinger, C. Fernandes, G. Formenti, I. Bista, P. R. Berg, C. Bleidorn, A. Bombarely, A. Crottini, G. R. Gallo, J. A. Godoy, S. Jentoft, J. Malukiewicz, A. Mouton, R. A. Oomen, S. Paez, P. J. Palsbøll, C. Pampoulie, M. J. Ruiz-Ĺopez, S. Secomandi, H. Svardal, C. Theofanopoulou, J. de Vries, A.-M. Waldvogel, G. Zhang, E. D. Jarvis, M. Balint, C. Ciofi, R. M. Waterhouse, C. J. Mazzoni, J. Hoglund, S. A. Aghayan, T. S. Alioto, I. Almudi, N. Alvarez, P. C. Alves, I. R. Amorim do Rosario, A. Antunes, P. Arribas, P. Baldrian, G. Bertorelle, A. Bohne, A. Bonisoli-Alquati, L. L. Bostjaňcíc, B. Boussau, C. M. Breton, E. Buzan, P. F. Campos, C. Carreras, L. F. C. Castro, L. J. Chueca, F. Čiampor, E. Conti, R. Cook-Deegan, D. Croll, M. V. Cunha, F. Delsuc, A. B. Dennis, D. Dimitrov, R. Faria, A. Favre, O. D. Fedrigo, R. Ferńandez, G. F. Ficetola, J.-F. Flot, T. Gabaldon, D. R. Agius, A. M. Giani, M. T. P. Gilbert, T. Grebenc, K. Guschanski, R. Guyot, B. Hausdorf, O. Hawlitschek, P. D. Heintzman, B. Heinze, M. Hiller, M. Husemann, A. Iannucci, I. Irisarri, K. S. Jakobsen, P. Klinga, A. Kloch, C. F. Kratochwil, H. Kusche, K. K. S. Layton, J. A. Leonard, E. Lerat, G. Liti, T. Manousaki, T. Marques-Bonet, P. Matos-Maravi, M. Matschiner, F. Maumus, A. M. McCartney, S. Meiri, J. Melo-Ferreira, X. Mengual, M. T. Monaghan, M. Montagna, R. W. Mys-lajek, M. T. Neiber, V. Nicolas, M. Novo, P. Ozretíc, F. Palero, L. P^arvulescu, M. Pascual, O. S. Paulo, M. Pavlek, C. Pegueroles, L. Pellissier, G. Pesole, C. R. Primmer, A. Riesgo, L. Rüber, D. Rubolini, D. Salvi, O. Seehausen, M. Seidel, B. Studer, S. Theodoridis, M. Thines, L. Urban, A. Vasemagi, A. Vella, N. Vella, S. C. Vernes, C. Vernesi, D. R. Vieites, C. W. Wheat, G. Worheide, Y. Wurm, and G. Zammit. How genomics can help biodiversity conservation. Trends in Genetics, 39(7):545–559, July 2023. ISSN 0168-9525. doi: 10.1016/j.tig.2023.01.005.

H. Wickham. ggplot2: Elegant Graphics for Data Analysis. Springer-Verlag New York, 2016. ISBN 978-3-319-24277-4. URL https://ggplot2.tidyverse.org.

C. Wiuf and J. Hein. Recombination as a point process along sequences. Theor Popul Biol, 55 (3):248–259, June 1999. ISSN 0040-5809. doi: 10.1006/tpbi.1998.1403.

