## Supplementary File S1 for "On the estimation of genome-average recombination rates": ResultsNotebook.html

##### Julien Y. Dutheil

#### 2023-05-23

### Ideal scenario

Read the data:

```
dat.cf <- read.csv("Results_ConstantPopSize_FlatRecombination.csv", row.names = 1)
```

First we compare SMC methods with others:

```
dat1 <- subset(dat.cf, GcProp == 0 & Method %in% c("MSMC1", "MSMC5", "iSMC1", "iSMC5", "LDhat5", "LDhat50", "Pyrho5", "Pyrho50", "heRhoCO1", "heRhoCO5", "heRhoCO50", "heRhoTotal1", "heRhoTotal5", "heRhoTotal50"))
dat1$Method <- factor(
  dat1$Method,
  levels = c("MSMC1", "MSMC5", "iSMC1", "iSMC5", "LDhat5", "LDhat50", "Pyrho5", "Pyrho50", "heRhoCO1", "heRhoCO5", "heRhoCO50", "heRhoTotal1", "heRhoTotal5", "heRhoTotal50"),
  labels = c("MSMC", "MSMC", "iSMC", "iSMC", "LDhat", "LDhat", "Pyrho", "Pyrho", "heRho", "heRho", "heRho", "heRho (total)", "heRho (total)", "heRho (total)")
  )
recs <- unique(dat1$SimRec)
```

Plot:

```
p <- ggplot(dat1, aes(x = SimRec, y = Rho)) +
  geom_point(shape = 1) + 
  facet_grid(SampleSize~Method, as.table = FALSE) +
  geom_abline(linetype = "dashed", aes(
    slope = ifelse((Method %in% c("MSMC", "iSMC") & SampleSize == 50) | (Method %in% c("LDhat", "Pyrho") & SampleSize == 1), 0, 1),
    intercept = 0)) +
  scale_x_continuous(trans = "sqrt", 
                     breaks = recs, 
                     labels = scales::label_number()) +
  scale_y_continuous(trans = "sqrt", 
                     breaks = recs, 
                     labels = scales::label_number()) +
  theme_pubr(x.text.angle = 45) +
  theme(panel.grid.major = element_line(linetype = "dotted")) +
  theme(panel.border = element_rect(fill = NA)) +
  xlab(expression("Simulated "*rho)) +
  ylab(expression("Inferred "*rho))
p
```

```
## Warning: Removed 34 rows containing missing values (`geom_point()`).
```

Some estimates are very high when the true rho is zero. We set a
limit to the y-axis:

```
p <- ggplot(dat1, aes(x = SimRec, y = Rho)) +
  geom_point(shape = 1) + 
  facet_grid(SampleSize~Method, as.table = FALSE) +
  geom_abline(linetype = "dashed", aes(
    slope = ifelse((Method %in% c("MSMC", "iSMC") & SampleSize == 50) | (Method %in% c("LDhat", "Pyrho") & SampleSize == 1), 0, 1),
    intercept = 0)) +
  scale_x_continuous(trans = "sqrt", 
                     breaks = recs, 
                     labels = scales::label_number(), 
                     limits = c(0, 0.03)) +
  scale_y_continuous(trans = "sqrt",
                     breaks = recs,
                     labels = scales::label_number(),
                     limits = c(0, 0.03)) +
  theme_pubr(x.text.angle = 45) +
  theme(panel.grid.major = element_line(linetype = "dotted")) +
  theme(panel.border = element_rect(fill = NA)) +
  xlab(expression("Simulated "*rho)) +
  ylab(expression("Inferred "*rho))
p
```

```
## Warning: Removed 103 rows containing missing values (`geom_point()`).
```

```
ggsave(p, filename = "Manuscript/Figure1.pdf", width = 10, height = 6)
```

```
## Warning: Removed 103 rows containing missing values (`geom_point()`).
```

#### Check demography

```
dat.demo <- read.csv("Results_ConstantPopSize_FlatRecombination_Demography.csv", row.names = 1)
dat.demo1 <- subset(dat.demo, GcProp == 0 & Method == "MSMC")
options("scipen" = 1)
dat.demo1$RecF <- factor(dat.demo1$SimRec, levels = recs, labels = paste0("rho == ", recs))

p.demo1 <- ggplot(dat.demo1, aes(x = Start, y = Ne, group = Replicate)) +
  geom_step(alpha = 0.3) + 
  scale_x_log10() + scale_y_log10(limits = c(1e3, 1e11)) +
  xlab("Time (generations)") +
  facet_grid(SampleSize~RecF, labeller = label_parsed) +
  geom_hline(yintercept = 1e5, linetype = "dashed") +
  theme_pubr(x.text.angle = 45) +
  theme(panel.grid.major = element_line(linetype = "dotted")) +
  theme(panel.border = element_rect(fill = NA))
p.demo1
```

```
## Warning: Transformation introduced infinite values in continuous x-axis
```

```
## Warning: Removed 3 rows containing missing values (`geom_step()`).
```

#### Try with less categories

```
dat.demo2 <- subset(dat.demo, GcProp == 0 & Method == "MSMC_16t")
options("scipen" = 1)
dat.demo2$RecF <- factor(dat.demo2$SimRec, levels = recs, labels = paste0("rho == ", recs))

p.demo2 <- ggplot(dat.demo2, aes(x = Start, y = Ne, group = Replicate)) +
  geom_step(alpha = 0.3) + 
  scale_x_log10() + scale_y_log10(limits = c(1e3, 1e11)) +
  xlab("Time (generations)") +
  facet_grid(SampleSize~RecF, labeller = label_parsed) +
  geom_hline(yintercept = 1e5, linetype = "dashed") +
  theme_pubr(x.text.angle = 45) +
  theme(panel.grid.major = element_line(linetype = "dotted")) +
  theme(panel.border = element_rect(fill = NA))
p.demo2
```

```
## Warning: Transformation introduced infinite values in continuous x-axis
```

```
## Warning: Removed 7 rows containing missing values (`geom_step()`).
```

Make a figure with only 5 individuals for the manuscript:

```
dat.demo3 <- subset(dat.demo, GcProp == 0 & Method %in% c("MSMC", "MSMC_16t") & SampleSize == 5)

options("scipen" = 1)
dat.demo3$RecF <- factor(dat.demo3$SimRec, levels = recs, labels = paste0("rho == ", recs))
labels = c("MSMC" = "Default", "MSMC_16t" = "16 intervals")

# Convergence failures:
dat.fail <- dat.cf %>% 
  filter(GcProp == 0) %>% 
  filter(Method %in% c("MSMC5", "MSMC5.16t")) %>%
  group_by(Method, SimRec) %>%
  dplyr::summarise(n = sum(!is.na(Rho)))
```

```
## `summarise()` has grouped output by 'Method'. You can override using the
## `.groups` argument.
```

```
dat.fail$Method <- gsub(dat.fail$Method, pattern = "5.", replacement = "_", fixed = TRUE)
dat.fail$Method <- gsub(dat.fail$Method, pattern = "5", replacement = "", fixed = TRUE)
dat.fail$x <- 5e4
dat.fail$y <- 1e10
dat.fail$RecF <- factor(dat.fail$SimRec, levels = recs, labels = paste0("rho == ", recs))

p.demo3 <- ggplot(dat.demo3, aes(x = Start, y = Ne)) +
  geom_step(alpha = 0.3, aes(group = Replicate)) + 
  scale_x_log10() + scale_y_log10(limits = c(1e3, 1e11)) +
  xlab("Time (generations)") +
  geom_label(data = dat.fail, aes(x = x, y = y, label = paste0(n, "/10"))) +
  facet_grid(Method~RecF, labeller = labeller(.rows = labels, .cols = label_parsed)) +
  geom_hline(yintercept = 1e5, linetype = "dashed") +
  theme_pubr(x.text.angle = 45) +
  theme(panel.grid.major = element_line(linetype = "dotted")) +
  theme(panel.border = element_rect(fill = NA))
p.demo3
```

```
## Warning: Transformation introduced infinite values in continuous x-axis
```

```
## Warning: Removed 3 rows containing missing values (`geom_step()`).
```

```
ggsave(p.demo3, filename = "Manuscript/SupplementaryFigure1.pdf", width = 8, height = 5)
```

```
## Warning: Transformation introduced infinite values in continuous x-axis
## Removed 3 rows containing missing values (`geom_step()`).
```

Better with less categories. But does it improve the estimation of
rho?

#### Tweaking the MSMC optimization

```
dat2 <- subset(dat.cf, GcProp == 0 & Method %in% c("MSMC5", "MSMC5.16t", "MSMC5.40", "MSMC5.r4"))
dat2$Method <- factor(
  dat2$Method,
  levels = c("MSMC5", "MSMC5.16t", "MSMC5.40", "MSMC5.r4"),
  labels = c("Default", "16~intervals", "40~iterations", "rho/theta == 4")
  )
recs <- unique(dat2$SimRec)
```

```
p <- ggplot(dat2, aes(x = SimRec, y = Rho)) +
  geom_point(shape = 1) + 
  facet_wrap(~Method, labeller = label_parsed) +
  geom_abline(linetype = "dashed", aes(
    slope = 1,
    intercept = 0)) +
  scale_x_continuous(trans = "sqrt",
                     breaks = recs, 
                     labels = scales::label_number(),
                     limits = c(0, 0.03)) +
  scale_y_continuous(trans = "sqrt",
                     breaks = recs, 
                     labels = scales::label_number(),
                     limits = c(0, 0.03)) +
  theme_pubr(x.text.angle = 45) +
  theme(panel.grid.major = element_line(linetype = "dotted")) +
  theme(panel.border = element_rect(fill = NA)) +
  xlab(expression("Simulated "*rho)) +
  ylab(expression("Inferred "*rho))
p
```

```
## Warning: Removed 52 rows containing missing values (`geom_point()`).
```

```
ggsave(p, filename = "Manuscript/SupplementaryFigure2.pdf", width = 5, height = 5)
```

```
## Warning: Removed 52 rows containing missing values (`geom_point()`).
```

#### Effect of sample size

We try from 1 to 5 diploid:

```
dat3 <- subset(dat.cf, GcProp == 0 & Method %in% c("MSMC1", "MSMC2", "MSMC3", "MSMC4", "MSMC5",
                                                "iSMC1", "iSMC2", "iSMC3", "iSMC4", "iSMC5"))
dat3$Method <- substr(dat3$Method, start = 1, stop = 4)
recs <- unique(dat3$SimRec)
```

Plotting the results:

```
p <- ggplot(dat3, aes(x = SimRec, y = Rho)) +
  geom_point(shape = 1) + 
  facet_grid(Method~SampleSize, labeller = label_parsed, as.table = FALSE) +
  geom_abline(linetype = "dashed", aes(
    slope = 1,
    intercept = 0)) +
  scale_x_continuous(trans = "sqrt",
                     breaks = recs,
                     labels = scales::label_number()) +
  scale_y_continuous(trans = "sqrt", 
                     breaks = recs, 
                     labels = scales::label_number()) +
  theme_pubr(x.text.angle = 45) +
  theme(panel.grid.major = element_line(linetype = "dotted")) +
  theme(panel.border = element_rect(fill = NA)) +
  xlab(expression("Simulated "*rho)) +
  ylab(expression("Inferred "*rho))
p
```

```
## Warning: Removed 71 rows containing missing values (`geom_point()`).
```

We set the y-limit as iSMC estimates for rho = 0 with 1 individual
are very high:

```
p <- ggplot(dat3, aes(x = SimRec, y = Rho)) +
  geom_point(shape = 1) + 
  facet_grid(Method~SampleSize, labeller = label_parsed, as.table = FALSE) +
  geom_abline(linetype = "dashed", aes(
    slope = 1,
    intercept = 0)) +
  scale_x_continuous(trans = "sqrt",
                     breaks = recs,
                     labels = scales::label_number(),
                     limits = c(0, 0.03)) +
  scale_y_continuous(trans = "sqrt",
                     breaks = recs,
                     labels = scales::label_number(),
                     limits = c(0, 0.03)) +
  theme_pubr(x.text.angle = 45) +
  theme(panel.grid.major = element_line(linetype = "dotted")) +
  theme(panel.border = element_rect(fill = NA)) +
  xlab(expression("Simulated "*rho)) +
  ylab(expression("Inferred "*rho))
p
```

```
## Warning: Removed 91 rows containing missing values (`geom_point()`).
```

Virtually no effect, apart from when the true rho is zero.

### Heterogeneous recombination landscape

#### Autocorelation random map:

Read the data:

```
dat.ch <- read.csv("Results_ConstantPopSize_VariableRecombination.csv", row.names = 1)
```

Filter:

```
dat1 <- subset(dat.ch, GcProp == 0 & Method %in% c("MSMC1.16t", "MSMC5.16t", "iSMC1", "iSMC5", "rhoSMC1", "rhoSMC5", "rhoSMC1post", "rhoSMC5post", "LDhat5", "LDhat50", "Pyrho5", "Pyrho50", "heRhoCO1", "heRhoCO5", "heRhoCO50", "heRhoTotal1", "heRhoTotal5", "heRhoTotal50"))
dat1$Method <- factor(
  dat1$Method,
  levels = c("MSMC1.16t", "MSMC5.16t", "iSMC1", "iSMC5", "rhoSMC1", "rhoSMC5", "rhoSMC1post", "rhoSMC5post", "LDhat5", "LDhat50", "Pyrho5", "Pyrho50", "heRhoCO1", "heRhoCO5", "heRhoCO50", "heRhoTotal1", "heRhoTotal5", "heRhoTotal50"),
  labels = c("MSMC", "MSMC", "iSMC", "iSMC", "rhoSMC (MLE)", "rhoSMC (MLE)", "rhoSMC (post.)", "rhoSMC (post.)", "LDhat", "LDhat", "Pyrho", "Pyrho", "heRho", "heRho", "heRho", "heRho (total)", "heRho (total)", "heRho (total)")
  )
recs <- unique(dat1$SimRec)
```

Plot:

```
p <- ggplot(dat1, aes(x = SimRec, y = Rho)) +
  geom_point(shape = 1) + 
  facet_grid(SampleSize~Method, as.table = FALSE) +
  geom_abline(linetype = "dashed", aes(
    slope = ifelse((Method %in% c("MSMC", "iSMC") & SampleSize == 50) | (Method %in% c("LDhat", "Pyrho") & SampleSize == 1), 0, 1),
    intercept = 0)) +
  scale_x_continuous(trans = "sqrt", 
                     breaks = recs, 
                     labels = scales::label_number(),
                     limits = c(0, 0.03)) +
  scale_y_continuous(trans = "sqrt",
                     breaks = recs,
                     labels = scales::label_number(),
                     limits = c(0, 0.03)) +
  theme_pubr(x.text.angle = 45) +
  theme(panel.grid.major = element_line(linetype = "dotted")) +
  theme(panel.border = element_rect(fill = NA)) +
  xlab(expression("Simulated "*rho)) +
  ylab(expression("Inferred "*rho))
p
```

```
## Warning: Removed 35 rows containing missing values (`geom_point()`).
```

```
ggsave(p, filename = "Manuscript/Figure2.pdf", width = 13, height = 6)
```

```
## Warning: Removed 35 rows containing missing values (`geom_point()`).
```

Compare the variance between MLE and posterior estimates of
rhoSMC:

```
dat.test <- subset(dat.ch, GcProp == 0 & Method %in% c("rhoSMC1", "rhoSMC5", "rhoSMC1post", "rhoSMC5post"))
dat.test$SampleSize <- as.ordered(dat.test$SampleSize)
dat.test$Method <- factor(dat.test$Method, levels = c("rhoSMC1", "rhoSMC5", "rhoSMC1post", "rhoSMC5post"), labels = c("MLE", "MLE", "Posterior", "Posterior"))
dat.test = dat.test %>% 
  group_by(EmpRec, Method, SampleSize) %>%
  summarise(Variance = var(Rho))
```

```
## `summarise()` has grouped output by 'EmpRec', 'Method'. You can override using
## the `.groups` argument.
```

```
m <- step(lm(Variance~EmpRec*Method*SampleSize, dat.test))
```

```
## Start:  AIC=-417.59
## Variance ~ EmpRec * Method * SampleSize
## 
##                            Df  Sum of Sq        RSS     AIC
## - EmpRec:Method:SampleSize  1 2.1758e-13 2.7441e-11 -419.47
## <none>                                   2.7223e-11 -417.59
## 
## Step:  AIC=-419.47
## Variance ~ EmpRec + Method + SampleSize + EmpRec:Method + EmpRec:SampleSize + 
##     Method:SampleSize
## 
##                     Df  Sum of Sq        RSS     AIC
## - Method:SampleSize  1 6.0000e-14 2.7501e-11 -421.43
## - EmpRec:Method      1 2.6230e-12 3.0063e-11 -420.00
## <none>                            2.7441e-11 -419.47
## - EmpRec:SampleSize  1 1.1949e-10 1.4693e-10 -394.62
## 
## Step:  AIC=-421.43
## Variance ~ EmpRec + Method + SampleSize + EmpRec:Method + EmpRec:SampleSize
## 
##                     Df  Sum of Sq        RSS     AIC
## - EmpRec:Method      1 2.6230e-12 3.0123e-11 -421.97
## <none>                            2.7501e-11 -421.43
## - EmpRec:SampleSize  1 1.1949e-10 1.4699e-10 -396.61
## 
## Step:  AIC=-421.97
## Variance ~ EmpRec + Method + SampleSize + EmpRec:SampleSize
## 
##                     Df  Sum of Sq        RSS     AIC
## - Method             1 3.6000e-14 3.0159e-11 -423.95
## <none>                            3.0123e-11 -421.97
## - EmpRec:SampleSize  1 1.1949e-10 1.4962e-10 -398.33
## 
## Step:  AIC=-423.95
## Variance ~ EmpRec + SampleSize + EmpRec:SampleSize
## 
##                     Df  Sum of Sq        RSS     AIC
## <none>                            3.0159e-11 -423.95
## - EmpRec:SampleSize  1 1.1949e-10 1.4965e-10 -400.32
```

```
hist(resid(m))
```

```
plot(resid(m)~fitted(m))
```

```
summary(m)
```

```
## 
## Call:
## lm(formula = Variance ~ EmpRec + SampleSize + EmpRec:SampleSize, 
##     data = dat.test)
## 
## Residuals:
##        Min         1Q     Median         3Q        Max 
## -2.427e-06 -8.392e-07 -2.059e-07  9.792e-07  2.635e-06 
## 
## Coefficients:
##                       Estimate Std. Error t value Pr(>|t|)    
## (Intercept)         -1.471e-06  5.180e-07  -2.839   0.0149 *  
## EmpRec               5.820e-04  3.409e-05  17.071 8.78e-10 ***
## SampleSize.L         9.598e-07  7.326e-07   1.310   0.2147    
## EmpRec:SampleSize.L -3.324e-04  4.821e-05  -6.895 1.66e-05 ***
## ---
## Signif. codes:  0 '***' 0.001 '**' 0.01 '*' 0.05 '.' 0.1 ' ' 1
## 
## Residual standard error: 1.585e-06 on 12 degrees of freedom
## Multiple R-squared:  0.9674, Adjusted R-squared:  0.9592 
## F-statistic: 118.6 on 3 and 12 DF,  p-value: 3.493e-09
```

No effect of method.

#### Partial decode map:

Read the data:

```
dat.cd <- read.csv("Results_ConstantPopSize_VariableRecombinationDecode.csv", row.names = 1)
```

Filter:

```
dat1 <- subset(dat.cd, GcProp == 0 & Method %in% c("MSMC1.16t", "MSMC5.16t", "iSMC1", "iSMC5", "rhoSMC1", "rhoSMC5", "rhoSMC1post", "rhoSMC5post", "LDhat5", "LDhat50", "Pyrho5", "Pyrho50", "heRhoCO1", "heRhoCO5", "heRhoCO50", "heRhoTotal1", "heRhoTotal5", "heRhoTotal50"))
dat1$Method <- factor(
  dat1$Method,
  levels = c("MSMC1.16t", "MSMC5.16t", "iSMC1", "iSMC5", "rhoSMC1", "rhoSMC5", "rhoSMC1post", "rhoSMC5post", "LDhat5", "LDhat50", "Pyrho5", "Pyrho50", "heRhoCO1", "heRhoCO5", "heRhoCO50", "heRhoTotal1", "heRhoTotal5", "heRhoTotal50"),
  labels = c("MSMC", "MSMC", "iSMC", "iSMC", "rhoSMC (MLE)", "rhoSMC (MLE)", "rhoSMC (post.)", "rhoSMC (post.)", "LDhat", "LDhat", "Pyrho", "Pyrho", "heRho", "heRho", "heRho", "heRho (total)", "heRho (total)", "heRho (total)")
  )
recs <- unique(dat1$SimRec)
```

Plot:

```
p <- ggplot(dat1, aes(x = SimRec, y = Rho)) +
  geom_point(shape = 1) + 
  facet_grid(SampleSize~Method, as.table = FALSE) +
  geom_abline(linetype = "dashed", aes(
    slope = ifelse((Method %in% c("MSMC", "iSMC") & SampleSize == 50) | (Method %in% c("LDhat", "Pyrho") & SampleSize == 1), 0, 1),
    intercept = 0)) +
  scale_x_continuous(trans = "sqrt",
                     breaks = recs,
                     labels = scales::label_number(),
                     limits = c(0, 0.03)) +
  scale_y_continuous(trans = "sqrt", 
                     breaks = recs, 
                     labels = scales::label_number(),
                     limits = c(0, 0.03)) +
  theme_pubr(x.text.angle = 45) +
  theme(panel.grid.major = element_line(linetype = "dotted")) +
  theme(panel.border = element_rect(fill = NA)) +
  xlab(expression("Simulated "*rho)) +
  ylab(expression("Inferred "*rho))
p
```

```
## Warning: Removed 18 rows containing missing values (`geom_point()`).
```

```
ggsave(p, filename = "Manuscript/SupplementaryFigure4.pdf", width = 13, height = 6)
```

```
## Warning: Removed 18 rows containing missing values (`geom_point()`).
```

#### Performance of rhoSMC when the recombination landscape is flat:

Filter:

```
dat4 <- subset(dat.cf, GcProp == 0 & Method %in% c("rhoSMC1", "rhoSMC1post", "rhoSMC5", "rhoSMC5post"))
dat4$Method <- factor(
  dat4$Method,
  levels = c("rhoSMC1", "rhoSMC1post", "rhoSMC5", "rhoSMC5post"),
  labels = c("MLE", "Posterior", "MLE", "Posterior")
  )
recs <- unique(dat4$SimRec)
```

Plot:

```
p <- ggplot(dat4, aes(x = SimRec, y = Rho)) +
  geom_point(shape = 1) + 
  facet_grid(SampleSize~Method, as.table = FALSE) +
  geom_abline(linetype = "dashed") +
  scale_x_continuous(trans = "sqrt",
                     breaks = recs, 
                     labels = scales::label_number()) +
  scale_y_continuous(trans = "sqrt", 
                     breaks = recs, 
                     labels = scales::label_number()) +
  theme_pubr(x.text.angle = 45) +
  theme(panel.grid.major = element_line(linetype = "dotted")) +
  theme(panel.border = element_rect(fill = NA)) +
  xlab(expression("Simulated "*rho)) +
  ylab(expression("Inferred "*rho))
p
```

```
## Warning: Removed 1 rows containing missing values (`geom_point()`).
```

Estimates are very high when the true rate is 0 and only one
individual is used. We set an upper limit for plotting clarity:

```
p <- ggplot(dat4, aes(x = SimRec, y = Rho)) +
  geom_point(shape = 1) + 
  facet_grid(SampleSize~Method, as.table = FALSE) +
  geom_abline(linetype = "dashed") +
  scale_x_continuous(trans = "sqrt", 
                     breaks = recs, 
                     labels = scales::label_number(),
                     limits = c(0, 0.04)) +
  scale_y_continuous(trans = "sqrt",
                     breaks = recs, 
                     labels = scales::label_number(),
                     limits = c(0, 0.04)) +
  theme_pubr(x.text.angle = 45) +
  theme(panel.grid.major = element_line(linetype = "dotted")) +
  theme(panel.border = element_rect(fill = NA)) +
  xlab(expression("Simulated "*rho)) +
  ylab(expression("Inferred "*rho))
p
```

```
## Warning: Removed 13 rows containing missing values (`geom_point()`).
```

```
ggsave(p, filename = "Manuscript/SupplementaryFigure5.pdf", width = 5, height = 5)
```

```
## Warning: Removed 13 rows containing missing values (`geom_point()`).
```

Compute how many points are not displayes:

```
with (na.omit(dat4), table(Rho > 0.04, Method, SimRec))
```

```
## , , SimRec = 0
## 
##        Method
##         MLE Posterior
##   FALSE  14        14
##   TRUE    6         6
## 
## , , SimRec = 0.0006
## 
##        Method
##         MLE Posterior
##   FALSE  20        19
##   TRUE    0         0
## 
## , , SimRec = 0.003
## 
##        Method
##         MLE Posterior
##   FALSE  20        20
##   TRUE    0         0
## 
## , , SimRec = 0.006
## 
##        Method
##         MLE Posterior
##   FALSE  20        20
##   TRUE    0         0
## 
## , , SimRec = 0.03
## 
##        Method
##         MLE Posterior
##   FALSE  20        20
##   TRUE    0         0
```

Can a Likelihood ratio test (LRT) or AIC comparison allow to identify
the best model for inferring rho?

AIC:

```
dat.aic.flat <- read.csv("Results_ConstantPopSize_FlatRecombination_AIC.csv", row.names = 1)
dat.aic.var<- read.csv("Results_ConstantPopSize_VariableRecombination_AIC.csv", row.names = 1)
dat.aic <- rbind(dat.aic.flat, dat.aic.var)
dat.aic = dat.aic %>% pivot_wider(
  id_cols = c("SampleSize", "RecombinationLandscape", "DemographicModel", "SimRec", "GcProp", "GcTrack", "Replicate"),
  names_from = "Method",
  values_from = "AIC")
dat.aic$diffAIC <- with(dat.aic, rhoSMC - iSMC)
dat.aic$RecombinationLandscape <- factor(dat.aic$RecombinationLandscape, levels = c("FlatRecombination", "VariableRecombination"), labels = c("Homogeneous", "Heterogeneous"))
```

```
dat.aic1 <- subset(dat.aic, GcProp == 0)
p.aic <- ggplot(dat.aic1, aes(x = SimRec, y = diffAIC)) +
  geom_point() +
  scale_y_continuous(trans = "pseudo_log",
                     breaks = c(-1e6, -1000, -10, 0, 10)) +
  scale_x_continuous(trans = "sqrt",
                     breaks = recs,
                     labels = scales::label_number()) +
  facet_grid(SampleSize~RecombinationLandscape) +
  theme_pubr(x.text.angle = 45) +
  theme(panel.grid.major = element_line(linetype = "dotted")) +
  theme(panel.border = element_rect(fill = NA)) +
  xlab(expression("Simulated "*rho)) +
  ylab(expression("AIC difference"))
p.aic
```

When the recombination is 0, the AIC selection can go very wrong and
favor rhoSMC. This most likely reflect convergence issues. When the
recombination rate is too low on average, AIC favors the simple iSMC
model.

Effect of GC:

```
dat.aic2 <- subset(dat.aic, SimRec == 0.006)
p.aic.gc <- ggplot(dat.aic2, aes(x = GcProp, y = diffAIC)) +
  geom_point() +
  scale_y_continuous(trans = "pseudo_log", 
                     breaks = c(-1e6, -1000, -10, 0, 10)) +
  scale_x_continuous(breaks = c(0, 0.1, 0.5, 0.9, 1), 
                     labels = scales::label_percent()) +
  facet_grid(SampleSize~RecombinationLandscape) +
  theme_pubr(x.text.angle = 45) +
  theme(panel.grid.major = element_line(linetype = "dotted")) +
  theme(panel.border = element_rect(fill = NA)) +
  xlab(expression("GC proportion")) +
  ylab(expression("AIC difference"))
p.aic.gc
```

Very high amount of GC lead to a higher false rejection of H0.

LRT:

```
dat.lik.flat <- read.csv("Results_ConstantPopSize_FlatRecombination_Likelihood.csv", row.names = 1)
dat.lik.var <- read.csv("Results_ConstantPopSize_VariableRecombination_Likelihood.csv", row.names = 1)
dat.lik <- rbind(dat.lik.flat, dat.lik.var)
dat.lik = dat.lik %>% pivot_wider(
  id_cols = c("SampleSize", "RecombinationLandscape", "DemographicModel", "SimRec", "GcProp", "GcTrack", "Replicate"),
  names_from = "Method",
  values_from = "logLik")
dat.lik$diffLik <- 2*with(dat.lik, rhoSMC - iSMC)
dat.lik$LRT <- pchisq(dat.lik$diffLik, df = 2, lower.tail = FALSE)
dat.lik$LRT[dat.lik$LRT < 1e-100] <- 1e-100
dat.lik$RecombinationLandscape <- factor(dat.lik$RecombinationLandscape, levels = c("FlatRecombination", "VariableRecombination"), labels = c("Homogeneous", "Heterogeneous"))
```

```
dat.lik1 <- subset(dat.lik, GcProp == 0)
p.lrt <- ggplot(dat.lik1, aes(x = SimRec, y = LRT)) +
  geom_point() + 
  scale_y_log10() +
  scale_x_continuous(trans = "sqrt",
                     breaks = recs, 
                     labels = scales::label_number()) +
  facet_grid(SampleSize~RecombinationLandscape) +
  theme_pubr(x.text.angle = 45) +
  theme(panel.grid.major = element_line(linetype = "dotted")) +
  theme(panel.border = element_rect(fill = NA)) +
  xlab(expression("Simulated "*rho)) +
  ylab(expression("LRT P-value"))
p.lrt
```

Gene conversion:

```
dat.lik2 <- subset(dat.lik, SimRec == 0.006)
p.lrt.gc <- ggplot(dat.lik2, aes(x = GcProp, y = LRT)) +
  geom_point() + 
  scale_y_log10() +
  scale_x_continuous(breaks = c(0, 0.1, 0.5, 0.9, 1), 
                     labels = scales::label_percent()) +
  facet_grid(SampleSize~RecombinationLandscape) +
  theme_pubr(x.text.angle = 45) +
  theme(panel.grid.major = element_line(linetype = "dotted")) +
  theme(panel.border = element_rect(fill = NA)) +
  xlab(expression("GC proportion")) +
  ylab(expression("LRT P-value"))
p.lrt.gc
```

Consistent with AIC, but the LRT nominal threshold to consider should
be much lower than the usual 1%.

Summary figure:

```
p <- ggarrange(p.aic, p.lrt, p.aic.gc, p.lrt.gc, nrow = 2, ncol = 2, labels = c("A", "B", "C", "D"))
ggsave(p, filename = "Manuscript/SupplementaryFigure6.pdf", width = 12, height = 12)
```

#### Inferrence of rho in presence of recombination hotspots:

Read the data:

```
dat.chs <- read.csv("Results_ConstantPopSize_RecombinationHotspots.csv", row.names = 1)
```

Filter:

```
dat1 <- subset(dat.chs, Method %in% c("MSMC1.16t", "MSMC5.16t", "iSMC1", "iSMC5", "rhoSMC1", "rhoSMC5", "rhoSMC1post", "rhoSMC5post", "LDhat5", "LDhat50", "Pyrho5", "Pyrho50", "heRhoCO1", "heRhoCO5", "heRhoCO50", "heRhoTotal1", "heRhoTotal5", "heRhoTotal50"))
dat1$Method <- factor(
  dat1$Method,
  levels = c("MSMC1.16t", "MSMC5.16t", "iSMC1", "iSMC5", "rhoSMC1", "rhoSMC5", "rhoSMC1post", "rhoSMC5post", "LDhat5", "LDhat50", "Pyrho5", "Pyrho50", "heRhoCO1", "heRhoCO5", "heRhoCO50", "heRhoTotal1", "heRhoTotal5", "heRhoTotal50"),
  labels = c("MSMC", "MSMC", "iSMC", "iSMC", "rhoSMC (MLE)", "rhoSMC (MLE)", "rhoSMC (post.)", "rhoSMC (post.)", "LDhat", "LDhat", "Pyrho", "Pyrho", "heRho", "heRho", "heRho", "heRho (total)", "heRho (total)", "heRho (total)")
  )
recs <- unique(dat1$SimRec)
```

Plot:

```
p <- ggplot(dat1, aes(x = SimRec, y = Rho)) +
  geom_point(shape = 1) + 
  geom_point(shape = 2, aes(y = EmpRec)) + 
  facet_grid(SampleSize~Method, as.table = FALSE) +
  geom_abline(linetype = "dashed", aes(
    slope = ifelse((Method %in% c("MSMC", "iSMC") & SampleSize == 50) | (Method %in% c("LDhat", "Pyrho") & SampleSize == 1), 0, 1),
    intercept = 0)) +
  scale_x_continuous(trans = "sqrt",
                     breaks = recs,
                     labels = scales::label_number(), limits = c(0, 0.03)) +
  scale_y_continuous(trans = "sqrt", 
                     breaks = recs, 
                     labels = scales::label_number()) +
  theme_pubr(x.text.angle = 45) +
  theme(panel.grid.major = element_line(linetype = "dotted")) +
  theme(panel.border = element_rect(fill = NA)) +
  xlab(expression("Simulated "*rho)) +
  ylab(expression("Inferred "*rho))
p
```

```
## Warning: Removed 82 rows containing missing values (`geom_point()`).
```

```
ggsave(p, filename = "Manuscript/Figure4.pdf", width = 13, height = 6)
```

```
## Warning: Removed 82 rows containing missing values (`geom_point()`).
```

### Demography

#### Decreasing population size

Read the data:

```
dat.dh <- read.csv("Results_DecreasingPopSize_VariableRecombination.csv", row.names = 1)
```

Filter:

```
dat1 <- subset(dat.dh, GcProp == 0 & Method %in% c("MSMC1.16t", "MSMC5.16t", "iSMC1", "iSMC5", "rhoSMC1", "rhoSMC5", "rhoSMC1post", "rhoSMC5post", "LDhat5", "LDhat50", "Pyrho5", "Pyrho50", "heRhoCO1", "heRhoCO5", "heRhoCO50", "heRhoTotal1", "heRhoTotal5", "heRhoTotal50"))
dat1$Method <- factor(
  dat1$Method,
  levels = c("MSMC1.16t", "MSMC5.16t", "iSMC1", "iSMC5", "rhoSMC1", "rhoSMC5", "rhoSMC1post", "rhoSMC5post", "LDhat5", "LDhat50", "Pyrho5", "Pyrho50", "heRhoCO1", "heRhoCO5", "heRhoCO50", "heRhoTotal1", "heRhoTotal5", "heRhoTotal50"),
  labels = c("MSMC", "MSMC", "iSMC", "iSMC", "rhoSMC (MLE)", "rhoSMC (MLE)", "rhoSMC (post.)", "rhoSMC (post.)", "LDhat", "LDhat", "Pyrho", "Pyrho", "heRho", "heRho", "heRho", "heRho (total)", "heRho (total)", "heRho (total)")
  )
recs <- c(0, 0.0006, 0.003, 0.006, 0.03)
```

Plot:

```
p <- ggplot(dat1, aes(x = SimRec, y = Rho)) +
  geom_point(shape = 1) + 
  facet_grid(SampleSize~Method, as.table = FALSE) +
  geom_abline(linetype = "dashed", aes(
    slope = ifelse((Method %in% c("MSMC", "iSMC") & SampleSize == 50) | (Method %in% c("LDhat", "Pyrho") & SampleSize == 1), 0, 1),
    intercept = 0)) +
  scale_x_continuous(trans = "sqrt",
                     breaks = recs,
                     labels = scales::label_number(), limits = c(0, 0.03)) +
  scale_y_continuous(trans = "sqrt", 
                     breaks = recs, 
                     labels = scales::label_number(), limits = c(0, 0.08)) +
  theme_pubr(x.text.angle = 45) +
  theme(panel.grid.major = element_line(linetype = "dotted")) +
  theme(panel.border = element_rect(fill = NA)) +
  xlab(expression("Simulated "*rho)) +
  ylab(expression("Inferred "*rho))
p
```

```
## Warning: Removed 4 rows containing missing values (`geom_point()`).
```

```
ggsave(p, filename = "Manuscript/Figure3.pdf", width = 13, height = 6)
```

```
## Warning: Removed 4 rows containing missing values (`geom_point()`).
```

#### Increasing population size

Read the data:

```
dat.ih <- read.csv("Results_IncreasingPopSize_VariableRecombination.csv", row.names = 1)
```

Filter:

```
dat1 <- subset(dat.ih, GcProp == 0 & Method %in% c("MSMC1.16t", "MSMC5.16t", "iSMC1", "iSMC5", "rhoSMC1", "rhoSMC5", "rhoSMC1post", "rhoSMC5post", "LDhat5", "LDhat50", "Pyrho5", "Pyrho50", "heRhoCO1", "heRhoCO5", "heRhoCO50", "heRhoTotal1", "heRhoTotal5", "heRhoTotal50"))
dat1$Method <- factor(
  dat1$Method,
  levels = c("MSMC1.16t", "MSMC5.16t", "iSMC1", "iSMC5", "rhoSMC1", "rhoSMC5", "rhoSMC1post", "rhoSMC5post", "LDhat5", "LDhat50", "Pyrho5", "Pyrho50", "heRhoCO1", "heRhoCO5", "heRhoCO50", "heRhoTotal1", "heRhoTotal5", "heRhoTotal50"),
  labels = c("MSMC", "MSMC", "iSMC", "iSMC", "rhoSMC (MLE)", "rhoSMC (MLE)", "rhoSMC (post.)", "rhoSMC (post.)", "LDhat", "LDhat", "Pyrho", "Pyrho", "heRho", "heRho", "heRho", "heRho (total)", "heRho (total)", "heRho (total)")
  )
recs <- c(0, 0.0006, 0.003, 0.006, 0.03)/2 #Ne is roughly 50000 in this dataset.
```

Plot:

```
p1 <- ggplot(dat1, aes(x = SimRec, y = Rho)) +
  geom_point(shape = 1) + 
  facet_grid(SampleSize~Method, as.table = FALSE) +
  geom_abline(linetype = "dashed", aes(
    slope = ifelse((Method %in% c("MSMC", "iSMC") & SampleSize == 50) | (Method %in% c("LDhat", "Pyrho") & SampleSize == 1), 0, 1),
    intercept = 0)) +
  scale_x_continuous(trans = "sqrt",
                     breaks = recs,
                     labels = scales::label_number(), limits = c(0, 0.015)) +
  scale_y_continuous(trans = "sqrt", 
                     breaks = recs, 
                     labels = scales::label_number(), limits = c(0, 0.08)) +
  theme_pubr(x.text.angle = 45) +
  theme(panel.grid.major = element_line(linetype = "dotted")) +
  theme(panel.border = element_rect(fill = NA)) +
  xlab(expression("Simulated "*rho)) +
  ylab(expression("Inferred "*rho))
p1
```

```
## Warning: Removed 146 rows containing missing values (`geom_point()`).
```

```
dat.ih2 <- read.csv("Results_IncreasingPopSize2_VariableRecombination.csv", row.names = 1)
```

Filter:

```
dat2 <- subset(dat.ih2, GcProp == 0 & Method %in% c("MSMC1.16t", "MSMC5.16t", "iSMC1", "iSMC5", "rhoSMC1", "rhoSMC5", "rhoSMC1post", "rhoSMC5post", "LDhat5", "LDhat50", "Pyrho5", "Pyrho50", "heRhoCO1", "heRhoCO5", "heRhoCO50", "heRhoTotal1", "heRhoTotal5", "heRhoTotal50"))
dat2$Method <- factor(
  dat2$Method,
  levels = c("MSMC1.16t", "MSMC5.16t", "iSMC1", "iSMC5", "rhoSMC1", "rhoSMC5", "rhoSMC1post", "rhoSMC5post", "LDhat5", "LDhat50", "Pyrho5", "Pyrho50", "heRhoCO1", "heRhoCO5", "heRhoCO50", "heRhoTotal1", "heRhoTotal5", "heRhoTotal50"),
  labels = c("MSMC", "MSMC", "iSMC", "iSMC", "rhoSMC (MLE)", "rhoSMC (MLE)", "rhoSMC (post.)", "rhoSMC (post.)", "LDhat", "LDhat", "Pyrho", "Pyrho", "heRho", "heRho", "heRho", "heRho (total)", "heRho (total)", "heRho (total)")
  )
recs <- c(0, 0.0006, 0.003, 0.006, 0.03)/10 #Ne is roughly 1400 in this dataset.
```

Plot:

```
p2 <- ggplot(dat2, aes(x = SimRec, y = Rho)) +
  geom_point(shape = 1) + 
  facet_grid(SampleSize~Method, as.table = FALSE) +
  geom_abline(linetype = "dashed", aes(
    slope = ifelse((Method %in% c("MSMC", "iSMC") & SampleSize == 50) | (Method %in% c("LDhat", "Pyrho") & SampleSize == 1), 0, 1),
    intercept = 0)) +
  scale_x_continuous(trans = "sqrt",
                     breaks = recs,
                     labels = scales::label_number(), limits = c(0, 0.0007)) +
  scale_y_continuous(trans = "sqrt", 
                     breaks = recs, 
                     labels = scales::label_number(), limits = c(0, 0.08)) +
  theme_pubr(x.text.angle = 45) +
  theme(panel.grid.major = element_line(linetype = "dotted")) +
  theme(panel.border = element_rect(fill = NA)) +
  xlab(expression("Simulated "*rho)) +
  ylab(expression("Inferred "*rho))
p2
```

```
## Warning: Removed 58 rows containing missing values (`geom_point()`).
```

```
p <- ggarrange(p2, p1, nrow = 2, ncol = 1, labels = c("A", "B"))
```

```
## Warning: Removed 58 rows containing missing values (`geom_point()`).
```

```
## Warning: Removed 146 rows containing missing values (`geom_point()`).
```

```
ggsave(p, filename = "Manuscript/SupplementaryFigure7.pdf", width = 13, height = 12)
```

```
summary(dat.dh$Theta)
summary(dat.ih$Theta)
summary(dat.ih2$Theta)
```

### Biased gene conversion

Get data with variable amount of gene conversion:

```
methods <- c("MSMC1.16t", "MSMC5.16t", "iSMC1", "iSMC5", "rhoSMC1", "rhoSMC5", "rhoSMC1post", "rhoSMC5post", "LDhat5", "LDhat50", "Pyrho5", "Pyrho50", "heRhoCO1", "heRhoCO5", "heRhoCO50", "heRhoTotal1", "heRhoTotal5", "heRhoTotal50")
dat.gc.cf <- subset(dat.cf, SimRec == 0.006 & Method %in% methods)
dat.gc.ch <- subset(dat.ch, SimRec == 0.006 & Method %in% methods)
dat.gc.dh <- subset(dat.dh, SimRec == 0.006 & Method %in% methods)
dat.gc <- rbind(dat.gc.cf, dat.gc.ch, dat.gc.dh)
dat.gc$Method <- factor(
  dat.gc$Method,
  levels = methods,
  labels = c("MSMC", "MSMC", "iSMC", "iSMC", "rhoSMC", "rhoSMC", "rhoSMC (post)", "rhoSMC (post)", "LDhat", "LDhat", "Pyrho", "Pyrho", "heRho", "heRho", "heRho", "heRho (total)", "heRho (total)", "heRho (total)")
  )
dat.gc$RecombinationLandscape <- factor(
  dat.gc$RecombinationLandscape,
  levels = c("FlatRecombination", "VariableRecombination"),
  labels = c("Flat", "Variable"))
dat.gc$DemographicModel <- factor(
  dat.gc$DemographicModel,
  levels = c("ConstantPopSize", "DecreasingPopSize"),
  labels = c("Constant", "Decreasing"))
dat.gc1 <- subset(dat.gc, SampleSize %in% c(1, 5, 50))
```

Plot the results. For simplification, we only plot sample size of 5
for the main figure:

```
dat.gc2 <- subset(dat.gc1, SampleSize == 5)
p <- ggplot(dat.gc2, aes(x = GcProp, y = Rho/0.006)) +
  geom_point(shape = 1) + 
  facet_grid(RecombinationLandscape+DemographicModel~Method, as.table = TRUE) +
  geom_hline(linetype = "dashed", yintercept = 1) +
  geom_abline(linetype = "dashed", slope = -1, intercept = 1) +
  geom_abline(linetype = "dashed", slope = 1, intercept = 1) +
  scale_x_continuous(labels = scales::label_percent()) +
  scale_y_continuous(labels = scales::label_percent(),
                     limits = c(0,2)) +
  theme_pubr(x.text.angle = 45) +
  theme(panel.grid.major = element_line(linetype = "dotted")) +
  theme(panel.border = element_rect(fill = NA)) +
  xlab(expression("Proportion of gene conversion")) +
  ylab(expression("Inferred "*rho))
p
```

```
## Warning: Removed 77 rows containing missing values (`geom_point()`).
```

```
ggsave(p, filename = "Manuscript/Figure5.pdf", width = 12, height = 8)
```

```
## Warning: Removed 77 rows containing missing values (`geom_point()`).
```

With one individual only:

```
dat.gc3 <- subset(dat.gc1, SampleSize == 1)
p <- ggplot(dat.gc3, aes(x = GcProp, y = Rho/0.006)) +
  geom_point(shape = 1) + 
  facet_grid(RecombinationLandscape+DemographicModel~Method, as.table = TRUE) +
  geom_hline(linetype = "dashed", yintercept = 1) +
  geom_abline(linetype = "dashed", slope = -1, intercept = 1) +
  geom_abline(linetype = "dashed", slope = 1, intercept = 1) +
  scale_x_continuous(labels = scales::label_percent()) +
  scale_y_continuous(labels = scales::label_percent(),
                     limits = c(0,2)) +
  theme_pubr(x.text.angle = 45) +
  theme(panel.grid.major = element_line(linetype = "dotted")) +
  theme(panel.border = element_rect(fill = NA)) +
  xlab(expression("Proportion of gene conversion")) +
  ylab(expression("Inferred "*rho))
p
```

```
## Warning: Removed 124 rows containing missing values (`geom_point()`).
```

```
ggsave(p, filename = "Manuscript/SupplementaryFigure8.pdf", width = 12*6/8, height = 8)
```

```
## Warning: Removed 124 rows containing missing values (`geom_point()`).
```

With 50 individuals:

```
dat.gc4 <- subset(dat.gc1, SampleSize == 50)
p <- ggplot(dat.gc4, aes(x = GcProp, y = Rho/0.006)) +
  geom_point(shape = 1) + 
  facet_grid(RecombinationLandscape+DemographicModel~Method, as.table = TRUE) +
  geom_hline(linetype = "dashed", yintercept = 1) +
  geom_abline(linetype = "dashed", slope = -1, intercept = 1) +
  scale_x_continuous(labels = scales::label_percent()) +
  scale_y_continuous(labels = scales::label_percent(),
                     limits = c(0,2)) +
  theme_pubr(x.text.angle = 45) +
  theme(panel.grid.major = element_line(linetype = "dotted")) +
  theme(panel.border = element_rect(fill = NA)) +
  xlab(expression("Proportion of gene conversion")) +
  ylab(expression("Inferred "*rho))
p
```

```
## Warning: Removed 16 rows containing missing values (`geom_point()`).
```

```
ggsave(p, filename = "Manuscript/SupplementaryFigure9.pdf", width = 12*4/7, height = 8)
```

```
## Warning: Removed 16 rows containing missing values (`geom_point()`).
```

### Genetic diversity

Read the data:

```
dat.lh <- read.csv("Results_ConstantPopSize10000_VariableRecombination.csv", row.names = 1)
```

Filter:

```
dat1 <- subset(dat.lh, GcProp == 0 & Method %in% c("MSMC1.16t", "MSMC5.16t", "iSMC1", "iSMC5", "rhoSMC1", "rhoSMC5", "rhoSMC1post", "rhoSMC5post", "LDhat5", "LDhat50", "Pyrho5", "Pyrho50", "heRhoCO1", "heRhoCO5", "heRhoCO50", "heRhoTotal1", "heRhoTotal5", "heRhoTotal50"))
dat1$Method <- factor(
  dat1$Method,
  levels = c("MSMC1.16t", "MSMC5.16t", "iSMC1", "iSMC5", "rhoSMC1", "rhoSMC5", "rhoSMC1post", "rhoSMC5post", "LDhat5", "LDhat50", "Pyrho5", "Pyrho50", "heRhoCO1", "heRhoCO5", "heRhoCO50", "heRhoTotal1", "heRhoTotal5", "heRhoTotal50"),
  labels = c("MSMC", "MSMC", "iSMC", "iSMC", "rhoSMC (MLE)", "rhoSMC (MLE)", "rhoSMC (post.)", "rhoSMC (post.)", "LDhat", "LDhat", "Pyrho", "Pyrho", "heRho", "heRho", "heRho", "heRho (total)", "heRho (total)", "heRho (total)")
  )
recs <- unique(dat1$SimRec)
```

Plot:

```
p <- ggplot(dat1, aes(x = SimRec, y = Rho)) +
  geom_point(shape = 1) + 
  facet_grid(SampleSize~Method, as.table = FALSE) +
  geom_abline(linetype = "dashed", aes(
    slope = ifelse((Method %in% c("MSMC", "iSMC") & SampleSize == 50) | (Method %in% c("LDhat", "Pyrho") & SampleSize == 1), 0, 1),
    intercept = 0)) +
  scale_x_continuous(trans = "sqrt",
                     breaks = recs,
                     labels = scales::label_number(), limits = c(0, 0.003)) +
  scale_y_continuous(trans = "sqrt", 
                     breaks = recs, 
                     labels = scales::label_number(), limits = c(0, 0.008)) +
  theme_pubr(x.text.angle = 45) +
  theme(panel.grid.major = element_line(linetype = "dotted")) +
  theme(panel.border = element_rect(fill = NA)) +
  xlab(expression("Simulated "*rho)) +
  ylab(expression("Inferred "*rho))
p
```

```
## Warning: Removed 36 rows containing missing values (`geom_point()`).
```

```
ggsave(p, filename = "Manuscript/SupplementaryFigure3.pdf", width = 13, height = 6)
```

```
## Warning: Removed 36 rows containing missing values (`geom_point()`).
```
